# Regulation of sex differences in innate immunity by sex-determining gene *transformer* in *Drosophila melanogaster*

**DOI:** 10.1101/2024.05.28.596272

**Authors:** MD Mursalin Khan, Rita M. Graze

**Affiliations:** Department of Biological Sciences, Auburn University, Alabama, USA

**Keywords:** *transformer*, *tra-*mutant, *tra-*overexpression, sex determination pathway, sex dimorphism, sex-biased gene expression, Toll, IMD, innate immunity, pseudomale, pseudofemale, intersex

## Abstract

Sex dimorphism is one of the key features of dioecious organisms, which shapes almost all the aspects of life history traits from early development to cessation of life. However, the specific regulatory differences between males and females that underlie sex dimorphism are underexplored for most quantitative traits, including immune sex dimorphism. Complex patterns of variation, for example, with respect to numbers and types of contributing alleles, dynamic changes in gene expression, and context dependency, have created challenges in understanding the mechanistic basis of sex differences in immunity. To investigate the regulatory basis of sex dimorphism in immunity, we focused on the sex determination hierarchy master switch, *transformer* (*tra*), in *Drosophila melanogaste*r. Two different perturbations of sex determination were examined, pseudomales (*tra-*mutant) and pseudofemales (*tra-*overexpression). Changes in gene expression patterns in response to the infection of *Providencia rettgeri*, an ex*tra*cellular gram-negative bacterium, were examined for each of these perturbations and for controls with wild-type sex determination phenotypes. To the best of our knowledge, this is the first study to report the immune response to bacterial infection in any insects with mutant or overexpressed *tra* genotypes. Survival and bacterial load were first characterized to assess the overall impact of *tra* on sex differences in immunity. To examine the regulatory role of the *tra* gene, sex differentially regulated genes were identified in comparisons of wound control to bacterial infected flies for males, females, and *tra-*mutant or *tra-*overexpression animals. The survival assay showed that in the absence of the female-specific isoform of *tra* (Tra^F^), *tra-*mutant XX animals showed significantly lower survival than wild-type females, and their survival rate aligned with males. The overexpression of Tra^F^ in the XY animals resulted in a surprising outcome, much higher survival of these animals relative to both males and females. The DEG analysis of genes with a significant interaction between sex and infection status identified 235 and 417 genes regulated upstream and downstream of *tra* in the sex hierarchy pathway, respectively. Interestingly, the GO enrichment analysis found 49 bacterial infection response-related biological processes enriched among genes regulated downstream of *tra*, including the Toll signaling pathway, and no enrichment for immune response-related categories among genes regulated upstream of *tra*. Further analysis of the genes and regulators of the Toll signaling pathway identified a significant regulatory role of *tra* or its downstream targets in signal detection, transduction, cellular response, and regulators of the pathway. Moreover, several genes regulated downstream of *tra* can regulate IMD pathways via the transcription factor NF-kB-Relish and some of its regulators, such as Diap2/IAP2 and Charon. Overall, the findings highlighted a strong potential role of *tra* to establish immune sex dimorphism in Drosophila.

## 1. Introduction

The sex determination pathway of *Drosophila* specifies sex differences, shaping almost all the aspects of its biology and life cycle (Fischer et al., 2015; Kraaijeveld et al., 2001; McDonald et al., 2021; Morrow, 2015; Wilkinson et al., 2022). While sex differences in easily noticeable attributes like size, shape, color, or structure have garnered widespread attention, it is essential to recognize that sexual dimorphism also encompasses less conspicuous traits such as immune responses and disease outcomes. These differences may not be immediately evident but nonetheless play a crucial role in reproduction and survival (Belmonte et al., 2020; Khodursky et al., 2020; Rai et al., 2023). Sex differences in immunity are detected in different life stages, for example, during reproductive phases, are under the influence of environmental stimuli, and can impact immunity throughout the animal or be localized to specific tissues (Deng & Jasper, 2016; Gal-Oz & Shay, 2022; Mackenzie et al., 2011; Wilkinson et al., 2022). These differences cannot be solely attributed to the regulation of sex hormones; they are also governed by other genetic and environmental influences (Brodin et al., 2015; Liston et al., 2021; Salz et al., 1989). Despite the prevalence and importance of sex differences in immunity, these differences have been largely overlooked even in model organisms and human studies, and therefore, whether the canonical sex determination pathway or one of its branches specifies these differences is unclear (Gal-Oz & Shay, 2022; Klein & Flanagan, 2016).

*Drosophila* has served as a valuable model organism for studying various aspects of biology, including immunity (Hoffmann, 2003). Sex dimorphism in the Drosophila immune response is well-established in multiple surveys of sex differences in immunity, revealing that males and females differ in their susceptibility to different pathogens, as well as in the expression of immune genes (Belmonte et al., 2020; Duneau, Kondolf, et al., 2017; Klein & Flanagan, 2016). For example, males and females may respond differently to viral, bacterial, fungal, and parasitic infections (Reviewed in (Belmonte et al., 2020; Klein & Flanagan, 2016)). The differences between males and females may reflect differing selective pressures due to their distinct reproductive roles and behaviors (McKean & Nunney, 2005; Scharf et al., 2013; Schwenke et al., 2016; Zuk & Stoehr, 2002).

The sex-dimorphic immune response is shaped by the complex interplay between immunity, mode of infection, reproductive status, and environmental conditions (McKean & Nunney, 2005, 2008; Nunn et al., 2009; Vincent & Sharp, 2014). The immune response can also vary depending on the stage of development of an organism (Jaillon et al., 2019; Klein & Flanagan, 2016; Millington & Rideout, 2018). Evolutionary processes, such as host-pathogen coevolution or sexual antagonism that shape allelic variation in genes underlying both immunity and sex dimorphism may add another layer of complexity (Khodursky et al., 2020; Lande, 1980; Morrow, 2015; Unckless et al., 2016). Thus, the genetics of sex differences in the immune response are likely to be complex, and this is reflected in the literature, where differences in candidate genes and pathways identified differ between studies (Belmonte et al., 2020).

Researchers have identified specific genes and signaling pathways that contribute to sexual dimorphism in the immune response of *Drosophila* (Belmonte et al., 2020; Duneau, Kondolf, et al., 2017; Vincent & Dionne, 2021). The work established that the classical bacterial response pathways, Toll and IMD, underlie sex dimorphism observed in *Drosophila melanogaster* (Duneau, Kondolf, et al., 2017; Vincent & Dionne, 2021). Research on sex dimorphism in the immune response is an expanding field; however, the complex molecular basis of this phenomenon remains poorly understood, and it is unclear which, if any, immunity-related pathways have the primary role in causing sex differences in the immune response and whether these immune pathways are themselves regulated by other genes such as sex determination genes, hormonal signaling genes, etc. (Graze et al., 2018; Schwenke & Lazzaro, 2017; Shepherd et al., 2021). The intricate crosstalk among evolutionarily conserved sex determination pathways and other physiological processes introduces further complexities in identifying the regulators of sex dimorphism in immunity. However, as sex determination is conserved, while sexually dimorphic traits are rapidly evolving, it might be possible to identify regulators that are commonly involved across species (Alejandro et al., 2022; Belote & Baker, 1987; Dudzic et al., 2019; Tafesh-Edwards & Eleftherianos, 2020). The first step will be to establish the top-level regulators in the *D. melanogaster* model.

Sex differences are likely to be regulated by the sex determination pathway in some way, but the regulatory connections between sex and immunity are unknown. The sex-determination process in *Drosophila* has long been understood to be initiated by the balance of factors on the X and the autosomes, although the dosage of X-linked factors may be the primary determinant according to more recent studies (Keyes et al., 1992; Salz & Erickson, 2010). In either model, the presence of different sex chromosome constitutions (XX or XY) leads to distinct expression patterns of the s*ex-lethal (sxl)* gene (Bell et al., 1991; Penalva & Sánchez, 2003). Sxl serves as a binary switch gene that regulates both sexual development and dosage compensation (Baker & Belote, 1983; Dahlsveen et al., 2006; Penalva & Sánchez, 2003). When two X chromosome copies are present, *sxl* expression is sufficient to trigger female-specific splicing of the downstream *transformer (tra)* pre-mRNA (Bell et al., 1991; Moschall et al., 2019). Consequently, functional Tra protein, a splicing factor, is produced exclusively in females (Keyes et al., 1992; McKeown et al., 1987; Salz & Erickson, 2010). Tra^F^ protein interacts with another splicing factor, Transformer-2 (Tra2), to bind with *doublesex (dsx)* pre-mRNA (Verhulst & van de Zande, 2015). In females, this interaction leads to the production of female-specific Dsx^F^ protein, which gives rise to female anatomical features (M. Arbeitman & Newell, 2016; Verhulst & van de Zande, 2015). In males, with only one X chromosome copy, an insufficient amount of X-linked signal elements (XSE) is produced, eventually leading to a lack of sufficient *sxl* expression; therefore, male-specific *tra* pre-mRNA contains an exon with a premature stop codon, thus generating nonfunctional Tra^M^ (Assis et al., 2012; Moschall et al., 2019). As a result, *doublesex (dsx)* and *fruitless (fru)* pre-mRNA undergo default splicing, yielding Dsx^M^ and Fru^M^ proteins (Assis et al., 2012; Salz & Erickson, 2010). The presence of Dsx^M^ specifies male anatomical features, and Fru^M^ works together with Dsx^M^ to develop male-specific neural development and mating behavior (Sato et al., 2019; Verhulst & van de Zande, 2015). In addition, the *transformer-2* (*tra-*2) gene plays a crucial role in governing somatic sexual differentiation in females and is essential for spermatogenesis in males (Belote & Baker, 1982; Goralski et al., 1989). In the context of the sex determination regulatory hierarchy, wild-type *tra-*2 function is necessary for the female-specific splicing of the pre-mRNA of the downstream gene, *doublesex (dsx)* (Belote & Baker, 1982; Goralski et al., 1989; Verhulst & van de Zande, 2015).

Collectively, these gene products play a pivotal role in shaping sex dimorphism, reproduction, and behavior (Salz et al., 1989). Previous research indicates that sex-determining transcription factors directly regulate the transcription of sex-dimorphic gene expression by binding to *cis*-regulatory elements of target genes, with some verified targets that are involved in both morphological and physiological traits (Chang et al., 2011; Keyes et al., 1992; Salz et al., 1989). Although numerous sex-dimorphic genes have been identified in insects, to the best of our knowledge, there is an absence of studies that investigate the transcriptomic profiles of sexually dimorphic genes when challenged with pathogenic infection in the context of mutations or overexpression of any sex-determination genes. Analyzing the differential changes in transcript levels upon pathogenic attack would greatly contribute to understanding the regulatory role of sex-determination genes in immune dimorphism.

The canonical sex-determination pathway is the key regulator of sex dimorphism in somatic cells (M. N. Arbeitman et al., 2016; Mank & Rideout, 2021). The disruption of the expression of the sex determination pathway genes such as *sxl, tra, dsx, fru,* and *msl-2 (male-specific lethal 2)* generate different phenotypes which may be described as intersex relative to wild-type females and males (Chang et al., 2011; Goldman & Arbeitman, 2007; Lyman et al., 1997; Rideout et al., 2015). In *Drosophila*, autosomal mutations in the *transformer (tra)* gene have been shown to transform chromosomal females (containing XX chromosomes) into phenotypic males (Fig. 1) (Wieschaus & Nöthiger, 1982). A *tra-*mutant XX animal, frequently referred to as a pseudomale in the literature, is an individual that has an XX genotype but exhibits male characteristics in terms of both morphology and behavior that are expressed in the absence of functional *transformer (tra)* or Tra^F^. The *tra-*mutant XX is also smaller than a wild-type female and does not develop functional ovaries (Rideout et al., 2015). The pseudomales (XX) do not exhibit *tra* expression, and consequently, there is no transcriptional influence from *sxl* that is propagated to *tra*, affecting the expression of *dsx* and *fru* genes, which ultimately leads to default splicing of *dsx* and *fru* to produce Dsx^M^ and Fru^M^ proteins. As a result, the *tra-*mutant animals develop male morphology and other male traits in a XX chromosomal fly. The presence of the XX will produce a sufficient amount of X-linked signal elements to activate *sxl*, but due to the absence of a functional *tra* allele, there is no impact of the *sxl* via the *sxl-tra-dsx* cascade (Fig. 1).

**Fig. 1:**
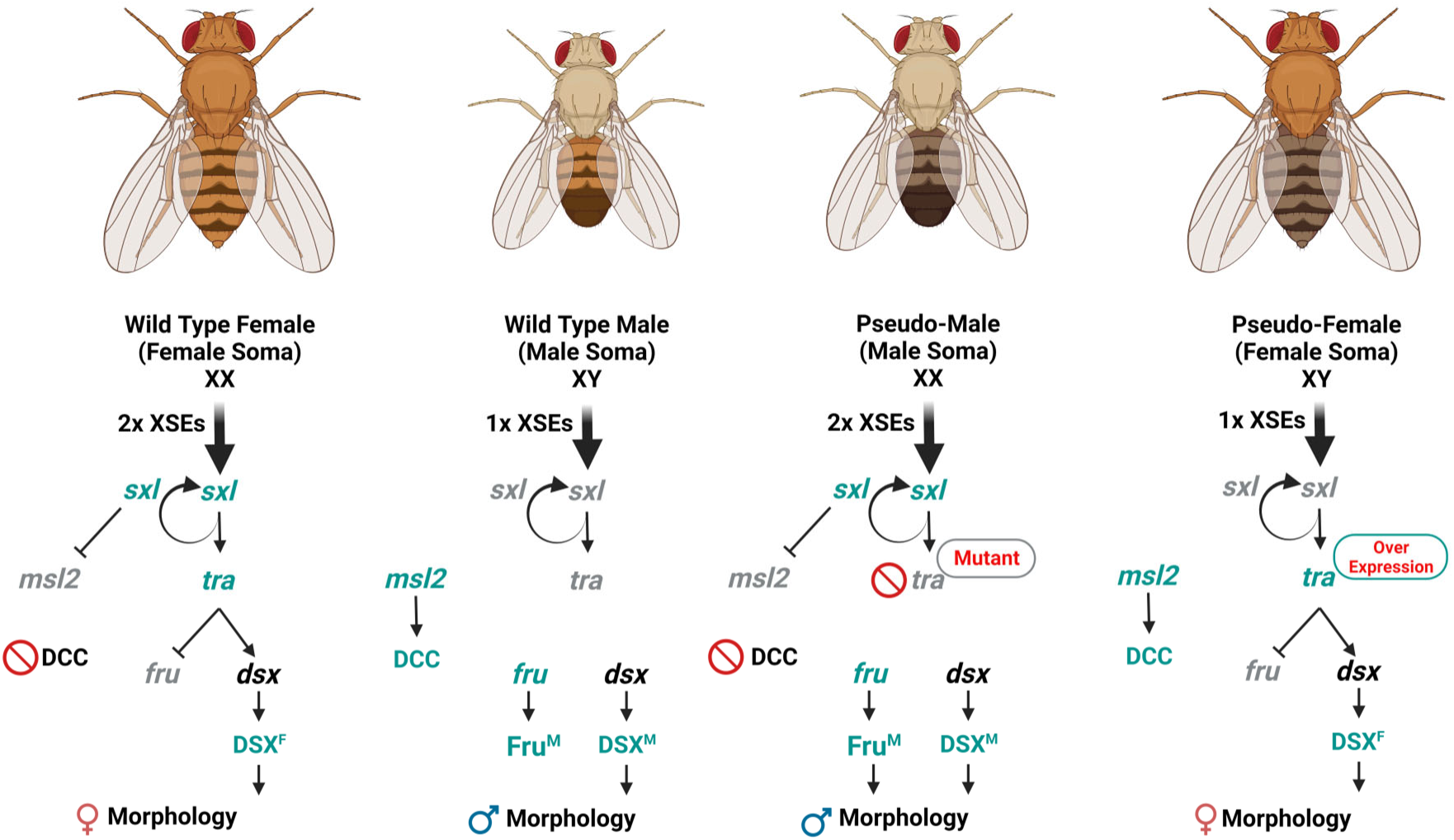
Drosophila sex determination hierarchy. The sex determination in the wild-type XX female, wild-type XY male, XX *tra-*mutant (pseudomale), and XY *tra-*overexpression (pseudofemale) animals. XSEs=X-linked signal elements, Sex-lethal (*sxl*), Transformer (*tra*), Doublesex (*dsx*), Fruitless gene (*fru*), Male-specific lethal 2 (*msl2*), Dosage compensation complex (DCC).

The overexpression of the *tra* gene in an XY animal also generates a distinct phenotype from wild-type females and males, which is commonly referred to as a pseudofemale (containing XY chromosomes) (Fig. 1). A *tra-*overexpressing XY individual exhibits female characteristics in terms of both morphology and behavior due to the overexpression of *tra* in an XY chromosomal fly (Rideout et al., 2015). The pseudofemale has an XY genotype; thus, the production of the X-linked signal elements is insufficient to produce enough of the Sxl protein. In the absence of sufficient Sxl protein, *tra-*overexpression is achieved by using the GAL4-UAS system, with a GAL4 driver and UAS-*tra* transgenic alleles used to overexpress *tra^F^*. Several GAL4 drivers have been used to overexpress *tra* (e.g., Actin5C-GAL4, Daughterless-GAL4), resulting in expression at a high level in the whole body or localized tissue expression (Rideout et al., 2015). For example, the Act5C-GAL4 driver-based expression of *tra* in an XY animal will result in ubiquitous expression throughout the body, in contrast to tissue-specific expression with fat body-specific *tra* overexpression (Rideout et al., 2015). The overexpression of ubiquitous Tra^F^ is not identical to the natural pattern of expression but results in the expression of female traits regulated downstream of *tra*. The *tra-*overexpression in the XY organism will lead to the expression of the female-specific *dsx*^F^ expression and inhibition of splicing to produce *fru*^M^, ultimately leading to female morphology and other characteristics (Fig. 1). For example, when a UAS-*tra^F^* transgene was expressed ubiquitously using the Daughterless-GAL4 driver, the *tra*-overexpressed flies exhibited a marked increase in body size relative to wild type males and females (Rideout et al., 2015).

In this study, the presence or absence of functional expression of core sex determination pathway gene *tra* was used to determine if there is a regulatory role of the sex determination pathway downstream of *tra* in immune sex dimorphism. To comprehensively characterize the role of *tra* in sex dimorphism in immunity in *Drosophila*, we conducted a comparative study using both *tra-*mutant and *tra-*overexpression strains and genetically matched controls with wild-type sex determination to observe the role of *tra* in the regulation of immune response against gram-negative extracellular bacteria *Providencia rettgeri*. This is the first study of bacterial infection in both *tra-*mutant and *tra-*overexpression animals reported in any species of insects, including Drosophila. Our study demonstrated that *tra* regulates sex differences in immunity. Through transcriptomic profiling, we identified specific immune response genes associated with these sex differences, as well as both the upstream and downstream genes with sex differences in expression in response to infection that are regulated by *tra*. We also find that ubiquitous overexpression of *tra* results in a large boost to the survival of infected flies relative to the wild type. The genes identified as regulated downstream of *tra* are putatively involved in sex dimorphism of the immune response and in the *tra* overexpression-mediated boost of the immune response.

## 2. Methods and Materials

### 2.1 Fly Husbandry

All flies were reared at 25°C with a 12:12 light: dark cycle on a cornmeal medium recipe (33 L H_2_O, 237 g agar, 825 g dried deactivated yeast, 1560 g cornmeal, 3300 g dextrose, 52.5 g Tegosept in 270 ml 95% ethanol and 60 ml propionic acid). The parental strains of both *tra-*mutant and *tra*-overexpression flies were obtained from the Bloomington Drosophila Stock Center (BDSC), Indiana University. Of note, the genetic backgrounds of these parental strains are different from each other, and in each case background, specific controls with wild-type sex determination were used for comparisons. The parental strains used to produce *tra* mutant animals were *tra*[1] (w[a]; *tra*[1]/TM2; BDSC-675(F)) and Df(3L)st-j7) (f[1]/Dp(1;Y)Bar[S]; Df(3L)st-j7, Ki[1]/TM6B, Tb[1]; BDSC-5416(M)), and for *tra* overexpression animals were Act5C-Gal4 (y[1] w[*]; P{w[+mC]=Act5C-GAL4}25FO1/CyO, y[+]; BDSC4414(F)) and UAS-*tra*.F (w[1118]; P{w[+mC]=UAS-*tra*.F}20J7; BDSC-4590(M)). All crosses were set up in vials with 15 females and 15 males, and progeny were collected as virgins and separated by sex. Female and male individuals were isolated in standard vials, and all experimental flies were unmated, adult, and aged 5-9 days prior to infection.

### 2.2 Sample collection

For the *tra-*mutant experiment, comparisons between females and males and between females and *tra-*mutant pseudomales collected from the same cross were used to categorize sex differences in expression in infected animals as regulated upstream or downstream of *tra*, following (Chang et al., 2011) (Fig. 1). The following genotypes were collected from the cross of *tra*[1] females to Df(3L)st-j7) males; XX females (w[a]/f[1]; TM2/TM6B), XY males (w[a]/ Dp(1;Y)Bar[S]; TM2/TM6B), and XX *tra-*mutant pseudomales (w[a]/ f[1]; *tra*[1]/Df(3L)st-j7, Ki[1]). In this study, we refer to the pseudomale (XX) or *tra*-mutant female (XX) as *tra-*mutant.

For the *tra-*overexpression experiment, differences between females and males and between males and animals with ubiquitous overexpression of *tra* collected from the same cross were examined to understand the role of *tra* in the increased effectiveness of the immune response, as observed in survival assays. The following genotypes were collected from the cross of Act5C-GAL4 females to UAS-*tra*.F males: XX females (y[1] w[*]/ w[1118]; P{w[+mC]=UAS-*tra*.F/CyO), XY males (y[1] w[*]/; P{w[+mC]=UAS-*tra*.F/CyO), and XY *tra-*overexpression pseudofemales (y[1] w[*]/; P{w[+mC]=UAS-*tra*.F/ P{w[+mC]=Act5C-GAL4). This study will refer to the pseudofemale (XY) or *tra*-overexpressed males (XY) as *tra-*overexpressed. Of note, the genetic backgrounds of the generated wild-type females and males differ between the *tra*-mutant and *tra*-overexpression strains; thus, the wild-type animals of each strain are not directly comparable, and background-specific controls were used in each experiment.

### 2.3 Experimental treatment

For the treatment, a strain of the gram-negative *Providencia rettgeri* species isolated from wild-caught *D. melanogaster* (*P. rettgeri* strain: Dmel) was used as an infectious agent, which is known as moderately virulent (Duneau, Kondolf, et al., 2017; Galac & Lazzaro, 2011; Juneja & Lazzaro, 2009). In this study, stationary phase *P. rettgeri* bacterial culture (OD600nm = 1.2) was used to inject into the thoracic region (soft spot near the wings) of the fly to introduce around 1200-1500 bacteria directly into the hemolymph using the needle prick method (Fig. 2) (Khalil et al., 2015). For control flies, sterile phosphate buffer solution (1x PBS) was injected as a wound-only control (Fig. 2) (Khalil et al., 2015). The flies were anesthetized by CO_2_ for less than 8 minutes during the injection. The fly vials were kept horizontal during recovery until the flies returned to normal activity levels.

**Fig. 2:**
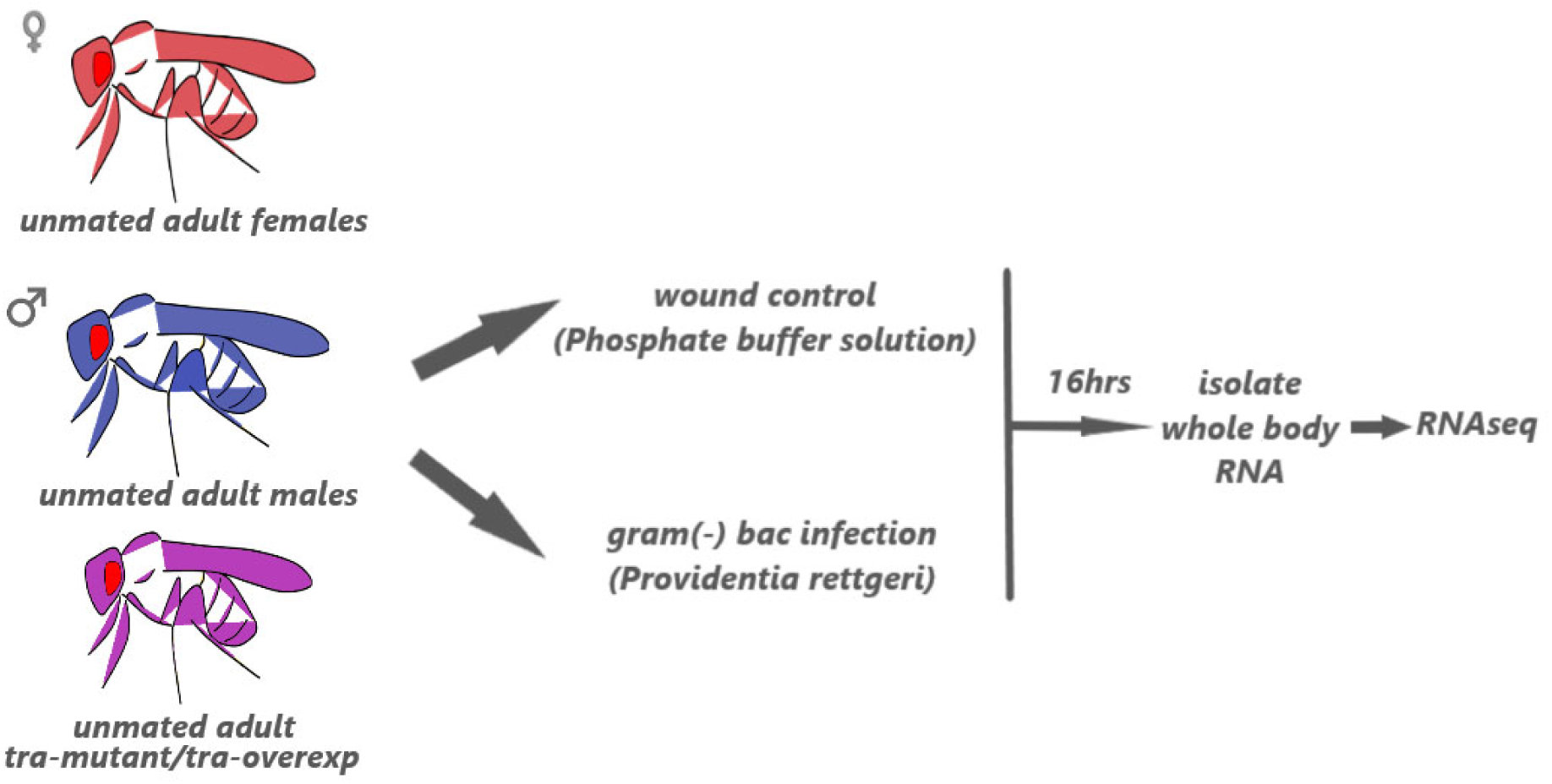
Experimental Design. Unmated adult flies were infected with *P. rettgeri* (PRET) by injection for the treatment and injected with phosphate buffer saline (PBS) for wound control. RNAseq was performed from the isolated RNA of the whole body at 16hpi in four replicates.

### 2.4 Bacterial culture preparation

Glycerol stocks of *P. rettgeri* were stored at −80°C prior to use. Stocks were used to streak LB-agar plates (Teknova, California) and incubated at 37°C for 12 hours to isolate single colonies. A single colony was then transferred to 5 mL Lysogeny broth (LB)-Broth (Teknova, California) and incubated at 37°C at 220 rotation per minute (rpm) for 9-10 hours. Stationary phase bacteria were centrifuged at 4000 rpm for 3 minutes to collect bacterial pellets. Pellets were washed with sterile 1x PBS using pipette aspiration (5 times) to remove residual LB Broth, and bacteria were recollected using the same centrifugal force. After five washes, the pellet was resuspended with sterile 1x PBS to achieve an OD600 of 1.2.

### 2.5 Survival rate

Four independent replicates of 5 to 9-day-old flies with 18 (*tra-*mutant) or 15 (*tra-*overexpressed) flies per sample were collected for each group and each treatment/control. Survival was scored at 2, 8, 16, 24, 36, and 48 hours (for *tra-*overexpression 72 hours) after bacterial injection. Flies were scored as dead if the fly was on its back and was not moving or could not turn over by itself (Duneau, Ferdy, et al., 2017). Scoring ceased when all the flies died in any experimental group or after 72hpi. Survival analysis was conducted in R using the package “survival” (Therneau, 2023). Kaplan-Meier survival curves were generated to visualize the survival proportion. The Cox regression was performed to determine the statistical difference among experimental groups (Therneau & Grambsch, 2000).

### 2.6 Bacterial load determination

A pool of three flies per replicate for each sample was homogenized in 200 µL 1X PBS using a sterile pestle. The homogenized sample (10 µL) was transferred to a 96-well plate and diluted with 90 µL of 1X PBS. From the first dilution (1/10), 10 µL of the solution was transferred successively, and serial dilution was carried out 8 times. Subsequently, 20 µL of the dilution 1:10^2,^ 1:10^4^, 1:10^6^, and 1:10^8^ were spread on LB agar plates to count CFU. Plates were incubated at 37 °C for 12-14 hours to count the CFU before gut microbes typically form colonies, which occurs at approximately 36 hours (Khalil et al., 2015). The bacterial load was determined at 8 and 16 hours after injections with four replicates of 5 to 9-day-old flies with 3 (*tra-*mutant) or 3 (*tra-*overexpressed) flies per sample (per pooled sample) following Duneau et al. (Duneau, Ferdy, et al., 2017). To identify significant differences for pairwise comparisons between groups, the linear model (CFU ∼ Groups) was fit (Sparks, 2017).

### 2.7 RNA isolation

Four independent replicates of 5 to 9-day-old flies with 18 (*tra-*mutant) or 15 (*tra-*overexpressed) flies per pool were collected for each genotype and group. Replicate pools of control and infected flies were flash-frozen in the liquid nitrogen at 16 hours post-injection (hpi). Only flies alive at 16hpi were used for RNA isolation. The frozen samples were stored at - 80°C prior to total RNA isolation. The 18 (*tra-*mutant) or 15 (*tra-*overexpressed) flies (per pooled sample) were homogenized in 700 µL of TRI-Reagent using a pellet pestle motor and RNase-Free pellet pestles in a 2 mL microcentrifuge tube. Total RNA was extracted using the Direct-zol RNA Miniprep Plus Kit (Zymo Research) according to the protocol of the manufacturer. The TURBO DNase-free kit (Thermo Fisher Scientific) was used to remove genomic DNA contamination following instructions of the manufacturer. The concentration was determined using Nanodrop 2.0 (Thermo Fisher Scientific), and the RNA integrity number (RIN) was measured using a tape-station system with an RNA screen-tape analysis kit (Agilent Technologies).

### 2.8 RNAseq library prep

RNAseq was performed by Novogene Corporation Inc., California. Messenger RNA was purified from total RNA using poly-T oligo-attached magnetic beads. After fragmentation, the first strand of cDNA was synthesized using random hexamer primers, followed by the second strand of cDNA synthesis. The library was ready after end repair, A-tailing, adapter ligation, size selection, amplification, and purification (Supp. File 1). Libraries were sequenced on a NovaSeq 6000 (Illumina), 150bp Paired-End reads, with ∼20 million reads per replicate sample (Supp. File 2).

### 2.9 Preprocessing and alignment

The sequenced reads were trimmed using Trimmomatic to remove the adaptors (Bolger et al., 2014). Quality assessment of raw data was performed by FastQC (Babraham Institute, University of Cambridge) (Anders, 2010). Reads were aligned against the genomic reference (FlyBase Releases: dmel_ r6.32 for *D. melanogaster*) using STAR (Gelbart et al., 1996; Gramates et al., 2022). The alignments were used to generate raw read counts for each gene and transcript in the reference genome annotation by using the “genomeDir”, “readFilesIn”, “quantMode”, and “outSAMtype” options in STAR (Dobin et al., 2013). Gene expression levels were then quantified using the “rsem-calculate-expression” option of RSEM (Li & Dewey, 2011).

### 2.10 Differential gene expression analysis

Differential expression analysis was conducted using an edgeR-limma-voom workflow (Ritchie et al., 2015; Robinson et al., 2010). Genes with fewer than 10 read counts (summed across samples within a group) were excluded from the analysis. Expression was estimated as the TMM normalized counts per gene and transcript for XX female, XY male, and XX *tra-*mutant (F, M, PM) and for XX female, XY male, and XY *tra-*overexpression (F, M, PF) animals, with infection by *P. rettgeri* (PRET) or PBS-only injected controls (PBS) separately. Voom variance modeling was implemented in limma, with a means model and comparisons between groups tested using contrasts (Law et al., 2014; Ritchie et al., 2015).

To verify expected differences between XX females, XY males, and XX *tra-*mutant pseudomales for the *tra-*mutant experiment and between XX, females, XY males, and XY *tra-*overexpression pseudofemales, each within treatment pairwise comparison was tested (Fig. 1). In the *tra-*mutant experiment, the effect of infection was examined using pairwise comparisons of treatment and control for each genotype: 1) female_pret = female_pbs, 2) male_pret = male_pbs, and 3) *tra-*mutant_pret = *tra-*mutant_pbs. To identify differences between genotypes in response to infection, the interaction of treatment by sex was tested with female_pret – female_pbs = male_pret – male_pbs, and the interaction of treatment by XX genotype was tested with female_pret – female_pbs = *tra-*mutant_pret – *tra-*mutant_pbs. The interaction of the treatment by male soma was also examined, with male_pret – male_pbs = *tra-*mutant_pret – *tra-*mutant_pbs. Expression differences regulated upstream and downstream of *tra* were categorized based on the following logic: FDR < 0.05 in interaction of treatment by sex and FDR > 0.05 in treatment by XX genotype for upstream of *tra* and FDR < 0.05 in interaction of treatment by sex and FDR < 0.05 in treatment by XX genotype for downstream of *tra* (Following Chang et al., 2011). For the *tra-*overexpression experiment, the main comparisons were female_pret = female_pbs, male_pret = male_pbs, and *tra-*overexp_pret = *tra-*overexp _pbs. To identify differences between genotypes in response to infection, the interaction of treatment by sex was tested with female_pret – female_pbs = male_pret – male_pbs, and the interaction of treatment by XY genotype was tested with male_pret – male_pbs = *tra-*overexp_pret – *tra-*overexp_pbs. The interaction of the treatment by female soma was also examined, with female_pret – female_pbs = *tra-*overexp_pret – *tra-*overexp_pbs. Unless otherwise mentioned, all the reported log2FoldChanges (log2FC) and interactions are significant at *p*-adj < 0.05.

### 2.11 Enrichment and pathway analyses

For enrichment tests, gene class heatmaps, and pathway analysis in *tra-*mutant and *tra-*overexpression experiments, the *D. melanogaster* gene and transcript ID were used (Supp. File 13). An FDR-adjusted *p*-value (*p*-adj) cutoff of 0.05 was used for all tests of differential gene expression. The “clusterProfiler” R program package was used for gene ontology (GO) based gene set enrichment analysis of differentially expressed genes (Wu et al., 2021). GO-Biological Process terms with adjusted *p*-value (FDR) less than 0.05 were considered significantly enriched. GO-BP annotations were performed separately for each contrast. The enrichment of DEGs in the Kyoto Encyclopedia of Genes and Genomes (KEGG) pathways was tested using the “clusterProfiler” package in the R (FDR < 0.05). Expression differences of DGEs for selected pathways were visualized using “Pathvisio” using the wiki pathways plugin. The wiki pathway WP3830 was used to generate the Toll-IMD-Jak/STAT gene list and pathways.

## 3. Results

Differences between males and females in immune responses are common in dioecious organisms, yet a full understanding of the regulatory genes and pathways involved remains elusive. However, sex-determination pathway genes are the central regulators of sex dimorphism in developmental and physiological processes (Belote & Baker, 1983; Mank & Rideout, 2021; Rideout et al., 2015; Salz et al., 1989). This study focuses on one of the key genes in the Drosophila sex determination pathway, *tra*. We investigate the role of *tra* in immune-related sex differences using *tra-*mutant and *tra-*overexpression flies (Fig. 1). The experimental groups were unmated wild-type females, wild-type males, *tra-*mutant (*pseudomales*, XX), and *tra-*overexpressed (*pseudofemale*, XY) (Fig. 1). The sex differences were assessed in control conditions and in animals infected with an extracellular gram-negative bacterium, *P. rettgeri*, using survival, bacterial load assays, and RNA-Seq-based gene expression (Fig. 2).

### 3.1 Survival assay of *tra-*mutant and *tra-*overexpression

An effective immune response against pathogenic infection is key to the survival of an organism (Fischer et al., 2015). Survival following *P. rettgeri* infection was assessed in *tra-*mutant animals over a period of 48 hours at 6-12-hour intervals (Fig. 3). During the infection initiation period (0-12hpi), no differences were observed among the female (XX), male (XY), and *tra-*mutant (XX) animals (Fig. 3A). The impact of the infection was apparent at 24hpi where females survived significantly higher than males and *tra-*mutant. At 36hpi, the female also survived highest among these three groups, and no differences between males and *tra-*mutant was detected. At 48hpi, no difference was identified in the survival of females, males, and *tra-*mutant as all the *P. rettgeri* infected samples died. Overall, the Kaplan Meier survival curve and Cox-regression model for females, males, and *tra-*mutant showed significant survival differences for the *P. rettgeri* infection. In the *tra-*mutant experiment, the females and males differed significantly in survival (*p-*value < 0.001). The *tra-*mutant also had significantly lower survival relative to females upon infection (*p-*value < 0.001). However, the comparison of the *tra-*mutant and males showed no significant difference in survival (*p-*value > 0.05). Both *tra* non-functional groups, males, and *tra-*mutants, showed similar survival curves for the *P. rettgeri* infection (Fig. 3A).

**Fig. 3:**
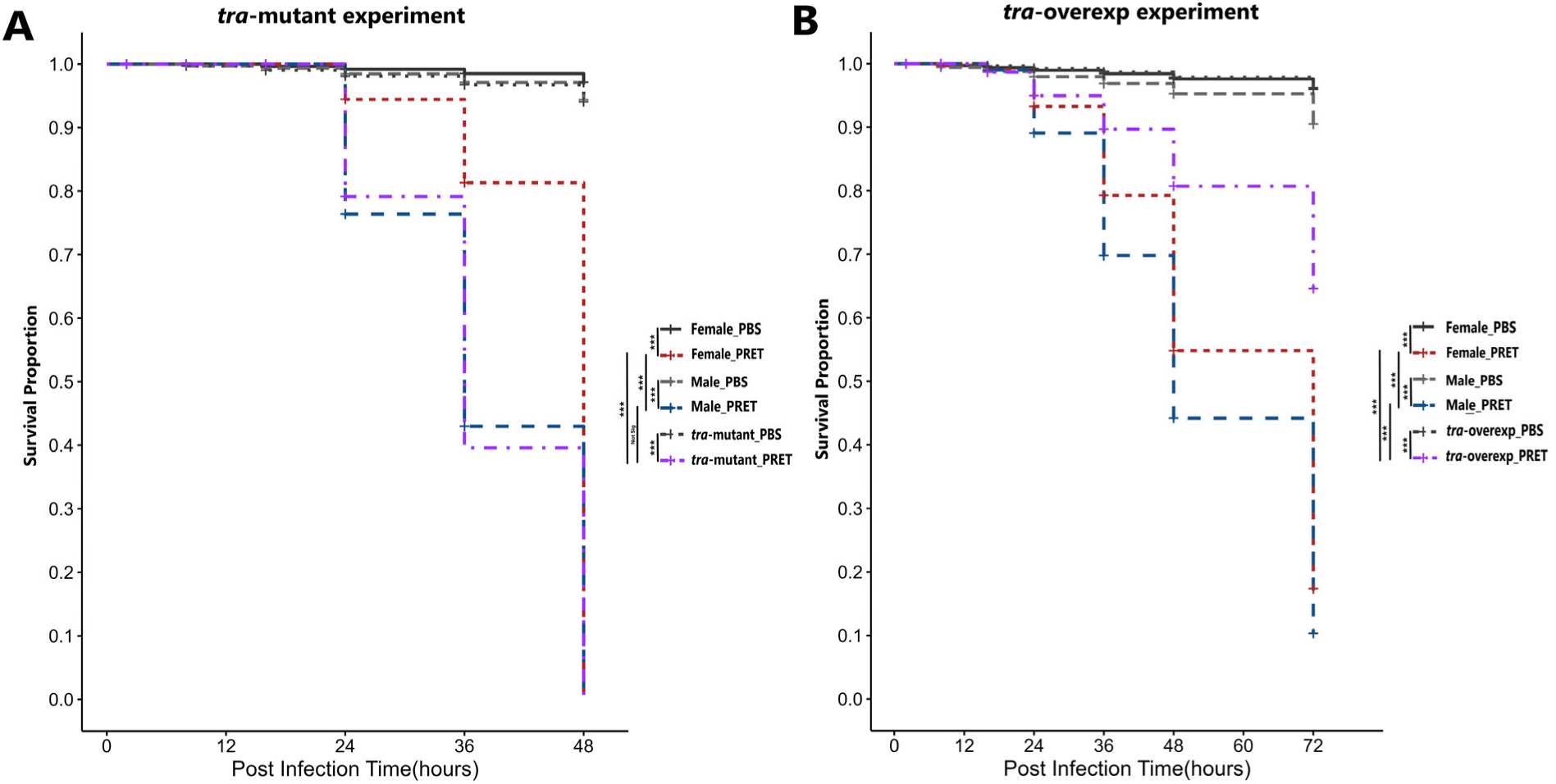
Survival assay for the *tra-*mutant and *tra-*overexpression experiments. **A.** Survival curve of wild-type XX females (Female), wild-type XY males (Male), and *tra-*mutant (pseudomale, PM) flies upon infection with *P. rettgeri* (PRET) and wound control (PBS). **B.** Survival curve of wild-type XX female (Female), wild-type XY male (Male), and XY *tra-*overexpression (*tra-*overexp, pseudofemale, PF) flies upon infection with *P. rettgeri* (PRET) and wound control (PBS). The survival curve was generated with the Kaplan–Meier estimator and the statistical analysis was performed with Cox regression. The asterisks (***) demons*tra*ted the *p-*values < 0.001, and Not-Sig represented No Significance. All the experimental flies (Female, Male, *tra-*mutant) of the *tra-mutant* experiment for PRET infection were identified dead at 48 hours.

Survival was similarly assayed in the *tra-*overexpressed flies over a period of 72hpi with *P. rettgeri* at 6-12-hour intervals (Fig. 3B). In the *tra-*overexpression experiment, the female (XX), male (XY), and *tra-*overexpressed (XY) animals showed trivial survival variation differences upon bacterial infection during the infection establishment period (0-12hpi) (Fig. 3B). The survival variation among the experimental groups was visible at 24hpi, where females survived significantly more than males (*p-*value < 0.001). However, the *tra-*overexpressed flies had significantly higher survival relative to both females and males at 24hpi. A consistent trend of survival was observed at 36hpi, where *tra-*overexpressed files showed the highest, and males showed the lowest survival. Finally, the 48- and 72-hpi time points showed a coherent pattern of survival for the females, males, and *tra-*overexpressed like 36hpi. Overall, the Kaplan Meier survival curve and Cox-regression model for females, males, and *tra-*overexpression animals showed significant survival differences for the *P. rettgeri* infection. In the *tra-*overexpressed experiment, the females survived significantly higher than males (*p-*value < 0.001), and the *tra-*overexpressed survived significantly higher than females (*p-*value < 0.001). Overall, the *tra-*overexpressed flies had the highest survival rates and much higher survival than either of the wild-type groups (Fig. 3).

### 3.2 Bacterial Load

The bacterial load determination at 8- and 16-hpi of the *tra-*mutant and *tra-*overexpression experiments revealed different patterns of changes in bacterial load. In the *tra-*mutant experiment, the 8- and 16hpi data of *P. rettgeri* infection showed no significant sex dimorphism among *tra*-mutant, females and males (Fig. 4A-B). For *tra-*overexpression, there was no dimorphism between females and males at 8hpi (Fig. 4C-D). At 16hpi, the females and *tra-*overexpressed flies demonstrated similar levels of bacterial load, whereas the bacterial load of males was significantly higher than both females (*p-*value < 0.001) and *tra-*overexpressed flies (*p-*value < 0.01) (Fig. 4D). Overall, the comparisons of bacterial load for *tra-*mutant experiment do not indicate that bacterial load underlies observed differences in survival rate; however, the *tra-*overexpression shows a similar pattern of the infection progression as identified in the survival assay, indicating that changes in bacteria load might have a relationship to observed differences in survival.

**Fig. 4:**
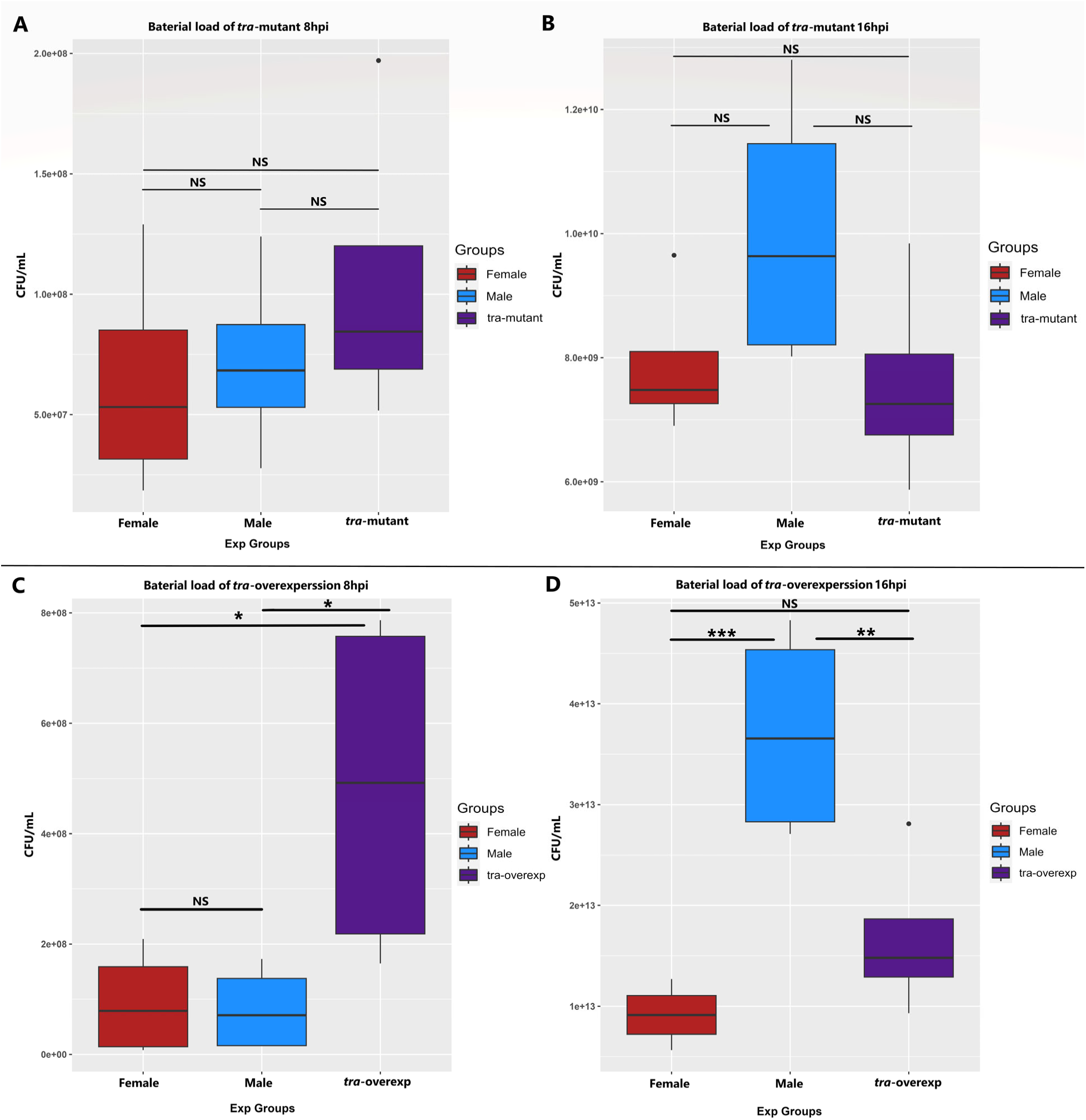
Bacterial load for the *tra* mutant and *tra-*overexpression upon *P. rettgeri* (PRET) infection. Determination of the bacterial load for the *tra-mutant* experiment of the wild-type females, wild-type males, and *tra-mutant* at 8hpi **(A)** and 16hpi **(B)**; for *tra-*overexpression experiment of the wild-type females, wild-type males and *tra-*overexpressed at 8hpi **(C)** and 16 hpi **(D)**. The asterisks (***) demons*tra*ted the *p-*values < 0.001, asterisks (**) demons*tra*ted the *p-*values < 0.01, asterisks (*) demons*tra*ted the *p-*values < 0.05, and NS represented No Significance. The wound control (PBS) data is not shown in Fig. due to the absence of bacterial growth in the 12-14-hour incubation period at 37°C.

### 3.3 Preprocessing and estimation of gene-level expression

Paired-end 150bp reads were generated using RNA Seq for each experimental group with 4 replicates. The total raw reads generated by RNA Seq was between 19.8 and 30.7 million (M) reads per sample, with an average of 22.7M for *tra-*mutant and 23.2M for *tra-*overexpression (Supp. File 2). The fastqc analysis showed that all 48 samples (24 for each experiment) had high quality throughout the read, and no trimming was performed except the removal of the adapter by the sequencing facility. The mean uniquely mapped read percentage for *tra-*mutant samples is 93.08%, and for *tra-*overexpression samples is 92.99%, which is typical for 20M reads of RNA Seq (Supp. File 2) (Conesa et al., 2016; Ma et al., 2019). Upon mapping, the raw read counts of 17869 genes in *tra-*mutant and *tra-*overexpression experiments were generated by STAR (Supp. File 3). Of the 17869 genes, 13,694 and 13,596 genes passed the detection cutoff for the *tra-*mutant and *tra-*overexpression experiments, respectively (Supp. File 4). After updating the gene annotation, the final analyzed gene counts were 13649 and 13553 for the *tra-*mutant and *tra-*overexpression experiments, respectively (Supp. File 5). Each experimental sample passed the quality control analysis prior to normalization and DEG analysis.

Normalization and DEG analysis were conducted using the edgeR and limma-voom packages in R (Ritchie et al., 2015; Robinson et al., 2010).

### 3.4 DEGs in *tra-*mutant and *tra-*overexpression

An analytical workflow comprising edgeR, limma, and voom was employed, incorporating a means model and contrasts for defined pairwise comparisons and interactions. This approach allowed for comparisons between wound control and infection treatments between specified experimental groups, with the goals of discerning regulatory changes in gene expression of infected flies across males, females, and XX *tra*-mutant or XY *tra*-overexpression animals (Ritchie et al., 2015; Robinson et al., 2010). The data was analyzed separately for *tra-*mutant and *tra-*overexpression experiments. Differences were classified as up or downregulated relative to wound control. The interaction of treatment by group was tested as a contrast to identify significant differences between the experimental groups within each experiment (Law et al., 2014; Ritchie et al., 2015). A significant treatment by group interaction term indicates that the groups differ in the way that the gene expression for a gene changes in response to the infection. Additional tests for interactions were included to classify changes in expression as up- or downstream of the *tra* gene.

Differential gene expression profiling of bacterial infection and wound control experimental groups identified differentially expressed genes for the *tra-*mutant and *tra-*overexpression experiments (Fig. 5, Supp. File 5). The expression profiles of the female, male, and *tra-*mutant were first analyzed under wound control only to understand patterns of differential expression resulting from differences in sex in the absence of bacterial infection (Fig. 5). In general, genes expected to differ between the sexes or to be perturbed in *tra*-mutant or *tra*-overexpression animals showed the expected patterns in these comparisons. For example, the female and *tra-*mutant comparison demonstrated that female (XX) upregulated yolk proteins (Yps) as expected, which is absent in the *tra-*mutants (XX) (Fig. 5A). On the other side, due to the absence of the *tra* in a XX female, there was higher expression of genes with strongly male-biased expression, such as accessory gland proteins (Acps), which is consistent with male-like gene expression in *tra-*mutants (Fig. 5A). In addition, comparing male to *tra-*mutant animals showed no differences in expression of accessory gland proteins (Acp), as both groups express these genes at a similar level (Fig. 5B). The absence of expression of female reproductive genes in the male and *tra-*mutant is consistent with the lack of development of ovaries in *tra-*mutant XX animals (Rideout et al., 2015).

**Fig. 5.**
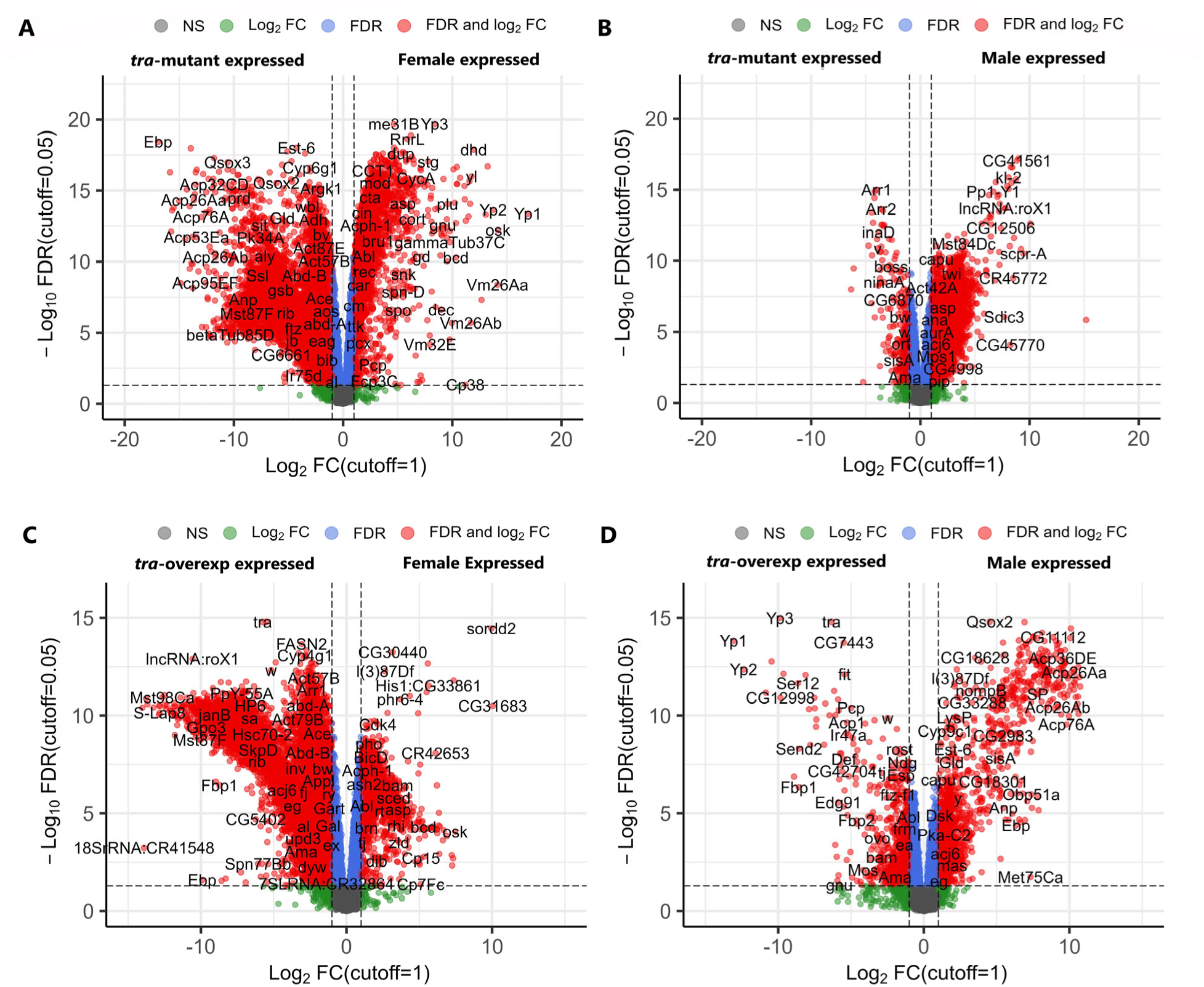
Volcano plots comparing each sex with the *tra-*mutant and *tra-*overexpression animals under control conditions (PBS injection). **A.** contrast between wild-type females and *tra-*mutants and **B.** contrast between wild-type males and *tra-*mutants; **C.** contrast between wild-type females and *tra-*overexpressed and **D.** contrast between wild-type males and *tra-*overexpressed are represented upon wound control (PBS). 13649 normalized genes with the symbol for *tra-*mutant and 13553 normalized genes with the symbol for *tra-*overexpression were used to represent DEGs with a cutoff: log2Fold Change ≥ 1; FDR < 0.05 (For details, see Supp. File 5).

In the *tra-*overexpression experiment, when comparing females with *tra-*overexpressed individuals, no significant upregulation was observed in female-biased proteins such as Yps. This lack of significance can be attributed to the fact that *Yp*s were expressed in both sets of flies at similar levels (Fig. 5C). However, when comparing males with *tra-*overexpressed individuals, a significant upregulation of *Yp*s was observed (Fig. 5D). Males also upregulated the genes expressed in male accessory glands, such as Sex-peptide (SP), when compared to XY *tra-*overexpression animals (Fig. 5D). Interestingly, the data revealed a significant upregulation of Andropin (Anp), an antibacterial peptide expressed in the male genital tract, in males due to the wound response (Fig. 5D). Finally, both comparisons showed that the XY *tra-*overexpressing flies upregulated the *tra* gene to significantly higher levels than in wild type females, and males (Fig. 5C-D).

Volcano plots of the two experimental groups show that males, females, XX *tra-*mutant, and XY *tra-*overexpressing animals broadly expressed key systemic immune and stress response genes at higher levels against gram-negative bacterial infection relative to the wound control animals (Fig. 6). The females and males of both experiments also showed sex dimorphism in the case of the up- and downregulation of genes in response to infection (Fig. 6).

**Fig. 6:**
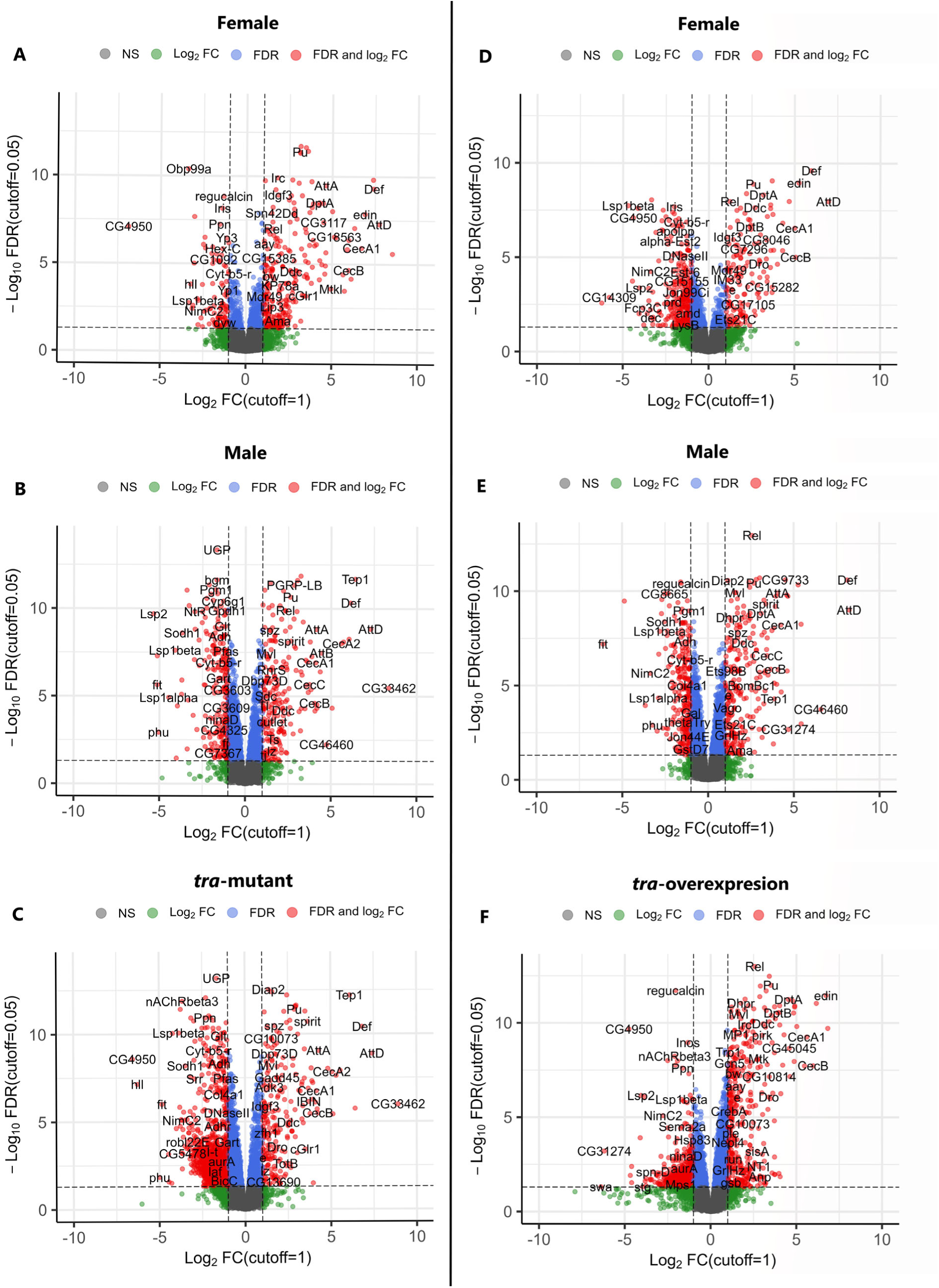
Volcano plots comparing the response to infection in each sex and in the *tra-*mutant and *tra-*overexpression animals. **A.** wild-type females, **B.** wild-type males, and **C.** *tra-*mutant males upon *P. rettgeri* (PRET) infection compared to wound control (PBS); **D.** wild-type females **E.** wild-type males and **F.** *tra-*overexpressed males upon *P. rettgeri* (PRET) infection compared to wound control (PBS). 13649 normalized genes with the symbol for *tra-*mutant and 13553 normalized genes with the symbol for tra-overexpression were used to represent DEGs with a cutoff: log2Fold Change ≥ 1; FDR < 0.05 (For details, see Supp. File 5).

In the *tra-*mutant study, females expressed several immune-related genes at significantly higher levels than the control, including AMPs like AttD, AttA, CecA1, CecB, DptA, and Mtkl (Fig. 6A). Females also upregulated the core transcription factor of the IMD pathway, NF-kB-Relish, to mount their immune response (Fig. 6A). Interestingly, females also upregulated negative regulatory proteins such as Serpin 42Dd, which suppress the Toll signaling pathway, which might indicate that females had initialized the suppression of immune response at the assayed time point (16hpi) (Fig. 6A). This pattern is consistent with suppression of the immune response when the infection is close to being controlled efficiently, compared to male and *tra-*mutant animals at the same timepoint (Khan2023_Unpublished_work_1, 2023; Khan2023_Unpublished_work_2, 2023). The females also downregulated Yolk proteins such as Yp1 and Yp2, which indicates plasticity in reproduction in response to infection, potentially due to trade-offs (Schwenke et al., 2016) (Fig. 6A). As previously reported, these results are consistent with independent observations of female and male expression differences under infection (Khan2023_Unpublished_work_1, 2023; Khan2023_Unpublished_work_2, 2023).

The male expressed several immune genes at a higher level compared to wound control, which was related to core bacterial infection response pathways, IMD and Toll (Fig. 6B). The genes include not only the effector molecules like AMPs but also signal receptors (PGRP-LB) and signal transducers of those pathways (*spz*, *spirit*) (Fig. 6B). In addition, males upregulated effector molecules of Jak-STAT (Hop-Stat92E) Pathways such as Turandot B (TotB) (Fig. 6B). Males also upregulated the Thioester-containing proteins (Tep1), which contributes to the humoral and cellular responses (Fig. 6B) (Dostálová et al., 2017). In the case of the downregulated genes in males, no negative immune regulators were detected, which is consistent with a higher deleterious impact of the infection in males than in females, as observed in the survival data (Fig. 6B, Fig. 3).

Lastly, in the absence of *tra* expression, the XX *tra-*mutant animals showed a more similar expression profile to males relative to females. The *tra*-mutant XX animals have a male soma due to the absence of Tra^F^; these pseudomales upregulate AMPs such as AttD, AttA, Def, CecA1, and CecA2 at a level similar to that of males (Fig. 6C). In addition, *tra-*mutants upregulate several signal transducers of the IMD and Toll pathways, such as *spz, spirit,* and *Diap2* (Fig. 6C). In concordance with the expected male expression pattern, XX *tra*-mutant animals also upregulate the effector molecules of the Jak-STAT (Hop-Stat92E) pathways, Turandot B (TotB) and Thioester-containing proteins (Tep1) (Fig. 6B-C). Overall, XY males and XX *tra-*mutant animals, which both lack TraF, show a similar expression profile (Fig. 6B-C). To further investigate the DEGs in detail, the genes that are regulated up- or downstream of *tra* were identified, and enrichment analysis was performed.

The *tra-*overexpression XY analysis also showed the expected sex differences between females and males (Fig. 6D-E). The females upregulated effector molecules related to IMD and Toll pathways, such as AttD, CecA1, CecB, Def, DptA, DptB, and Dro (Fig. 6D). The females also upregulated defense response-related genes like *immune molecule 33(IM33), edin*, and *Ets21C* (Fig. 6D). The expression profile of the males was distinct from that of females, consistent with different levels of infection-related stress conditions in males compared to females, as the survival and bacterial load data suggest (Fig. 3-4). The males upregulated IMD and Toll pathway-derived AMPs such as AttA, AttD, CecA1, CecB, CecC, Def, and DptA (Fig. 6E). In addition, the males induced the immune response by upregulating the *spz*, *spirit*, and Diap2 like signaling molecules of IMD and Toll pathways (Fig. 6E). Males also upregulated the Thioester-containing proteins (Tep1) and Toll-induced effector molecule Bomanin Bicipital 1 (BomBc1) (Fig. 6E).

Finally, the Actin5C promoter-based XY *tra-*overexpressing flies showed a diverging pattern of DEGs upon gram-negative infection compared to XY males. The *pseudofemales* expressed AMPs such as Anp, CecA1, CecB, Dro, DptA, DptB, and Mtk (Fig. 6F). The *tra-*overexpressed flies also uniquely upregulated a serine protease, Melanization Protease 1 (MP1), at a higher level, which plays an essential role in the melanization based immune responses (Fig. 6F). This pattern is explored further in the enrichment analysis for *tra-*overexpression experimental flies to gain insights into the unexpected large benefit to survival of the infection in *tra*-overexpressing animals.

### 3.5 Comparison of gene expression profiles in female, male, and *tra-*mutant/overexp

In the comparison within experiments, after normalization, 13694 genes were tested and classified as up- or downregulated in response to infection for *tra-*mutant experiment. The males downregulated 1139 genes compared to 267 downregulated by females in total; both sexes commonly downregulated 200 genes. The greater number of downregulated genes in males suggests a greater stress response in males upon infection (Fig. 7, Supp. File 6). The number of upregulated genes was also different between wild-type females and males. Overall, the significantly upregulated gene count for males is 922, and in females, 518, where both upregulated 323 genes in common (Fig. 7). The *tra-*mutant, which is an XX-*pseudomale*, had a unique set of DEGs relative to wild-type males and females. In total, the *tra-*mutant upregulated 921 genes and downregulated remarkably 3719 genes upon infection with *P. rettgeri*. The *tra-*mutant and males commonly upregulated 636 genes and downregulated 865 genes, whereas the *tra-*mutant and females commonly upregulated 337 genes and downregulated 214 genes. Overlap between all three, female, male, and *tra-*mutant, was not extensive, with 298 commonly upregulated genes and 193 downregulated genes (Fig. 7).

**Fig. 7:**
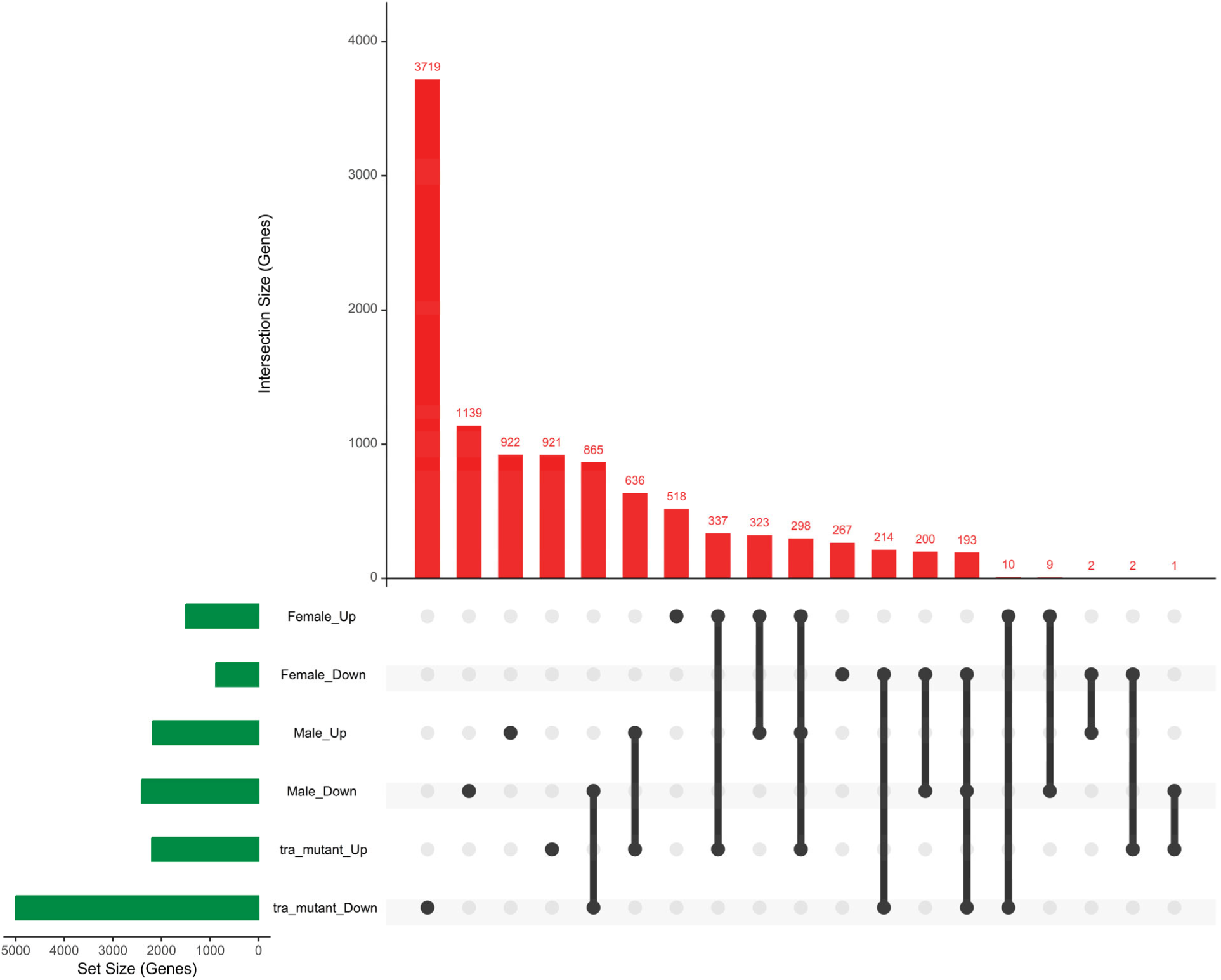
Up and downregulated gene counts for the *tra-*mutant experiment. **A.** Comparison across wild-type XX females, wild-type XY males, and the XX *tra-*mutant. The plots include synergistically and antagonistically expressed gene numbers for each group in the *tra-*mutant experiments. The figure shows an intersection-based representation (For details, see Supp. File 6).

After normalization, 13596 genes were tested in the *tra-*overexpression experiment. Overall, the males downregulated 1075 genes compared to 689 downregulated by females; both sexes commonly downregulated 451 genes, which could reflect higher stress levels in males as in the *tra* mutant experiment (Fig. 8, Supp. File 6). The number of upregulated genes was also different in both sexes for wild-type females and males, similar to the pattern observed in the *tra-*mutant experiment. In total, the significantly upregulated gene count for males is 939, and in females, it is 328, where both upregulated 223 genes in common (Fig. 8). The *tra-*overexpressing animals (XY-*pseudofemales)*, which had a much higher rate of survival, also differed greatly in patterns of gene expression, with a much greater number of genes upregulated (n = 1767) and downregulated (n = 1640). The XY *tra-*overexpression animals and males commonly upregulated 606 genes and downregulated 256 genes. The *tra-*overexpression animals and females commonly upregulated 219 genes and downregulated 146 genes. The female, male, and *tra-*overexpression commonly upregulated 198 genes and downregulated 128 genes (Fig. 8, Supp. File 6).

**Fig. 8:**
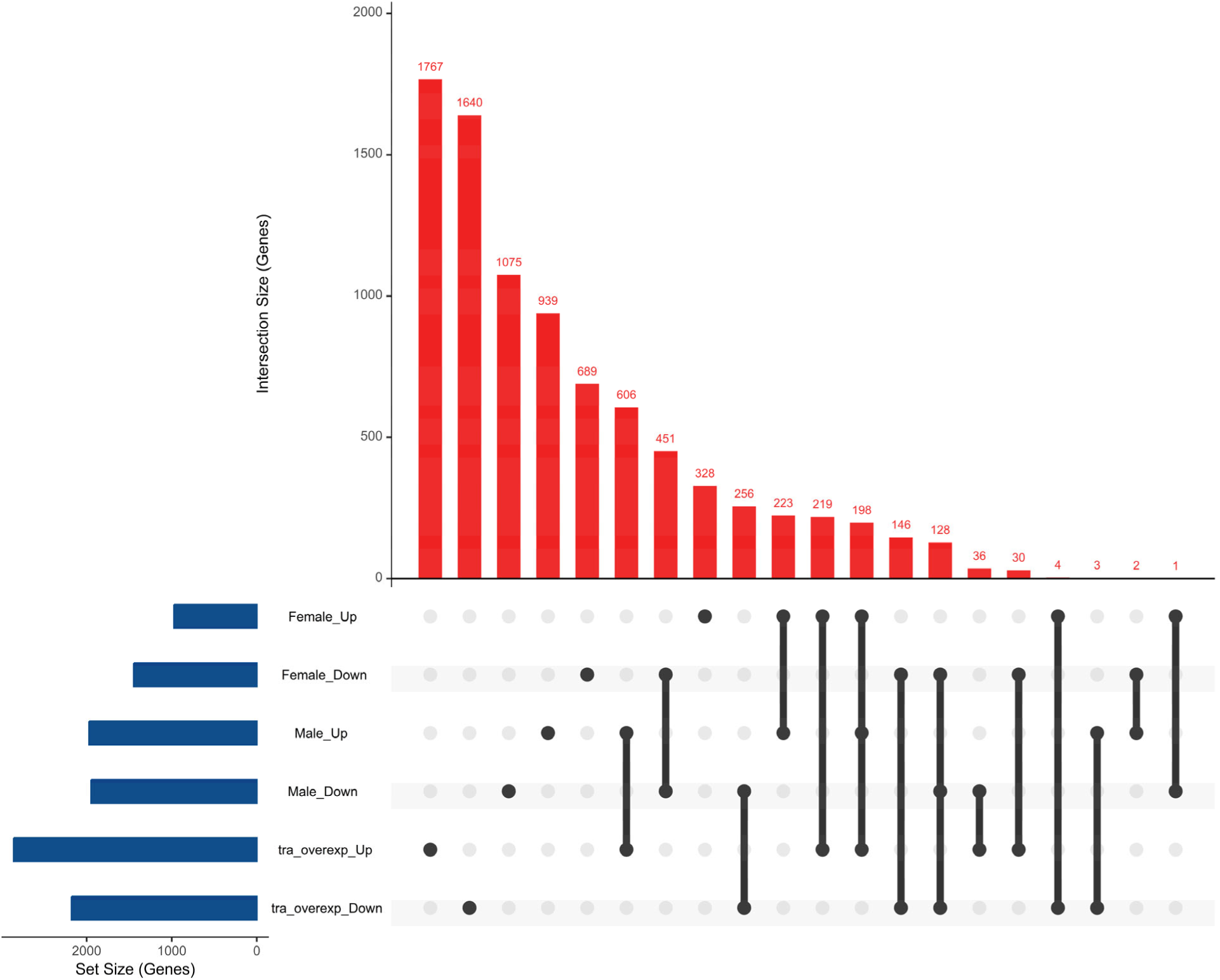
Up and down-regulated gene counts for the *tra-*overexpression experiment. **A.** Comparison across wild-type XX females, wild-type XY males, and XY *tra-*overexpression animals. The plots include synergistically and antagonistically expressed gene numbers for each group in the *tra-*mutant experiments. The figure uses an intersection-based representation (For details, see Supp. File 6).

### 3.6 Comparison of enriched GO terms in *tra*-mutant/overexpression experiments

Gene set enrichment analysis (GSEA) was conducted using the gene ontology biological process (GO-BP) for DEGs (FDR 0.05 and > 2Fold-Change (2FC)) for the *tra-*mutant and *tra-*overexpression experiments. In the experiment, the enriched gseGO-BP terms showed that females, males, and *tra-*mutants upregulated similar immune and defense response gene sets but downregulated distinct sets of genes during bacterial infection (Fig. 9, Supp. File 9). Among the upregulated DEGs (FDR < 0.05 and 2FC), 33 GO-BP terms were overrepresented in females. These terms were predominantly related to immunity. In contrast, there were 5 GO-BP terms overrepresented among downregulated genes, and these terms were associated with genes involved in metabolic processes. There were 66 GO-BP terms enriched among upregulated DEGs (FDR 0.05 and 2FC) in males, and 18 GO-BP terms overrepresented among downregulated DEGs at 16hpi in the *tra-*mutant experiment. Males also upregulated genes involved primarily in the immune response and positive regulators; however, there were some terms related to the negative regulation of metabolic processes and macromolecule metabolic processes. In the case of downregulated genes, males downregulated several metabolic processes, for example, lipid, carboxylic acid, and cellular carbohydrate. etc. The *tra-*mutant showed enrichment of 169 terms; interestingly, only one term was found to be overrepresented in the downregulated genes, mitochondrion organization, and the rest of the 168 terms corresponded to upregulated DEGs. Similar to the results for females and males, *tra-*mutant enriched terms were largely those related to immune response and regulation. However, the *tra-*mutant showed several unique enrichments related to development and morphogenesis (Fig. 9, Supp. File 9).

**Fig. 9:**
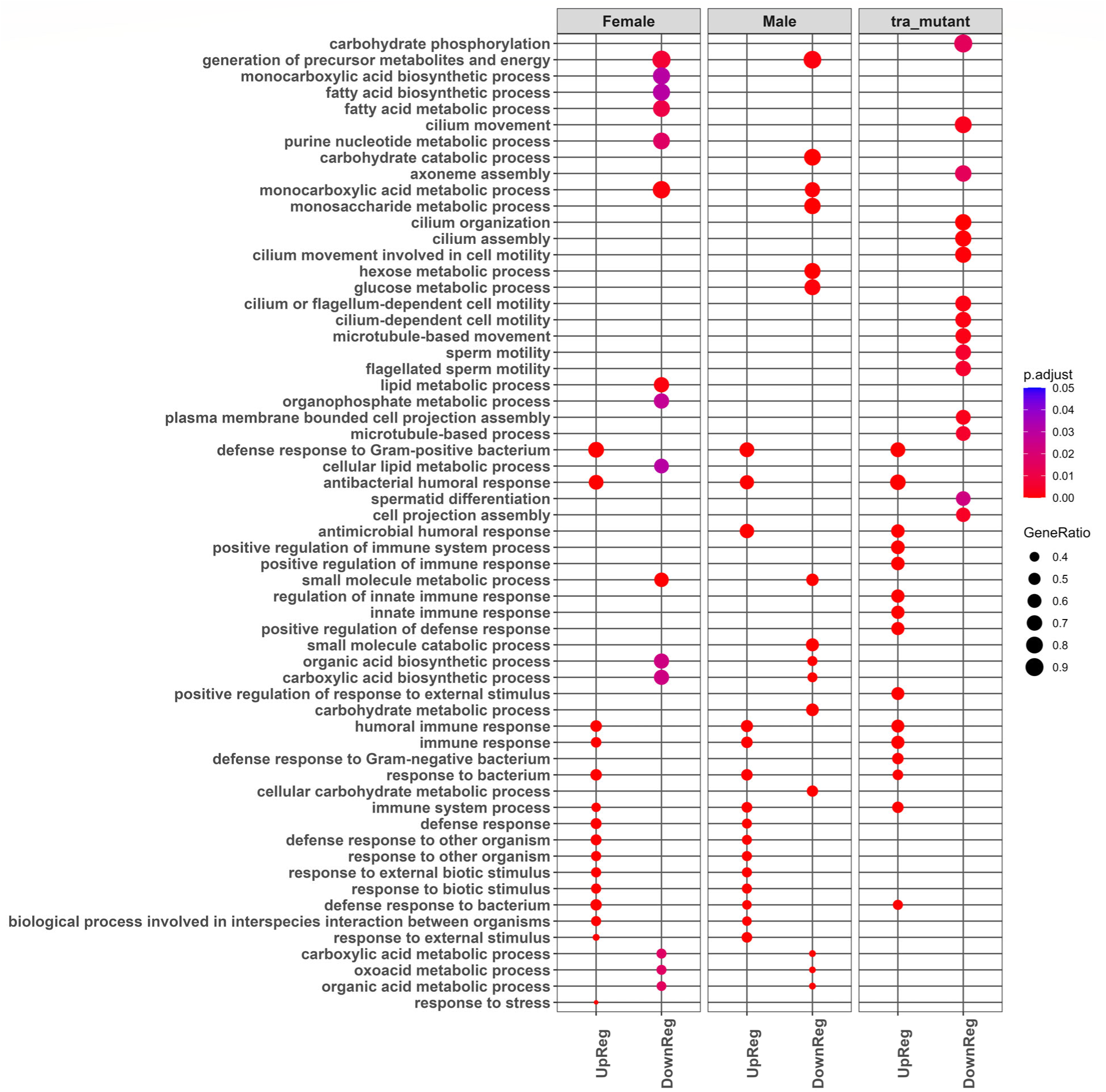
Top15 up and down-regulated Enriched Gene Ontology terms (gseGO: BP) in females, males, and *tra-*mutant upon *P. rettgeri* (PRET) infection (For details Supp. File 9). UpReg=Up Regulation, DownReg=Down Regulation.

In the *tra-*overexpression experiment, the enriched gseGO-BP terms showed that females, males, and *tra-*overexpression groups upregulated immune and defense response gene sets like the *tra-*mutant experiment, but they used different approaches to inhibit bacterial progression (Fig. 10, Supp. File 9). There were 39 categories overrepresented among upregulated DEGs (FDR 0.05 and 2FC) of females; among them, immune response and regulation-related terms were dominant. Among the 31 terms corresponding to downregulated genes, reproductive and developmental terms were frequent. Males upregulated 37 terms; interestingly, all of these were related to the immune response. The 9 downregulated terms of the males were connected to metabolic processes. Finally, the *tra-*overexpression showed a remarkable 399 enriched biological processes (FDR 0.05 and 2FC). Interestingly, out of these 399 terms, only 66 corresponded to upregulated genes, and the rest corresponded to downregulated genes. For upregulation, most of the gene sets were related to the immune response; however, the enriched regulation list showed some crucial negative regulatory terms such as negative regulation of response to biotic stimulus, negative regulation of response to external stimulus, and negative regulation of immune system process which might indicate the initiation of immune suppression in *tra-*overexpression flies. The majority of the enriched terms among downregulated genes were related to reproduction, development, and cell division. Although the upregulated enriched GO-BP terms were comparable to XX females, the 333 enriched GO-BP terms among downregulated genes were mostly distinct from those found in females (Fig. 10, Supp. File 9).

**Fig. 10:**
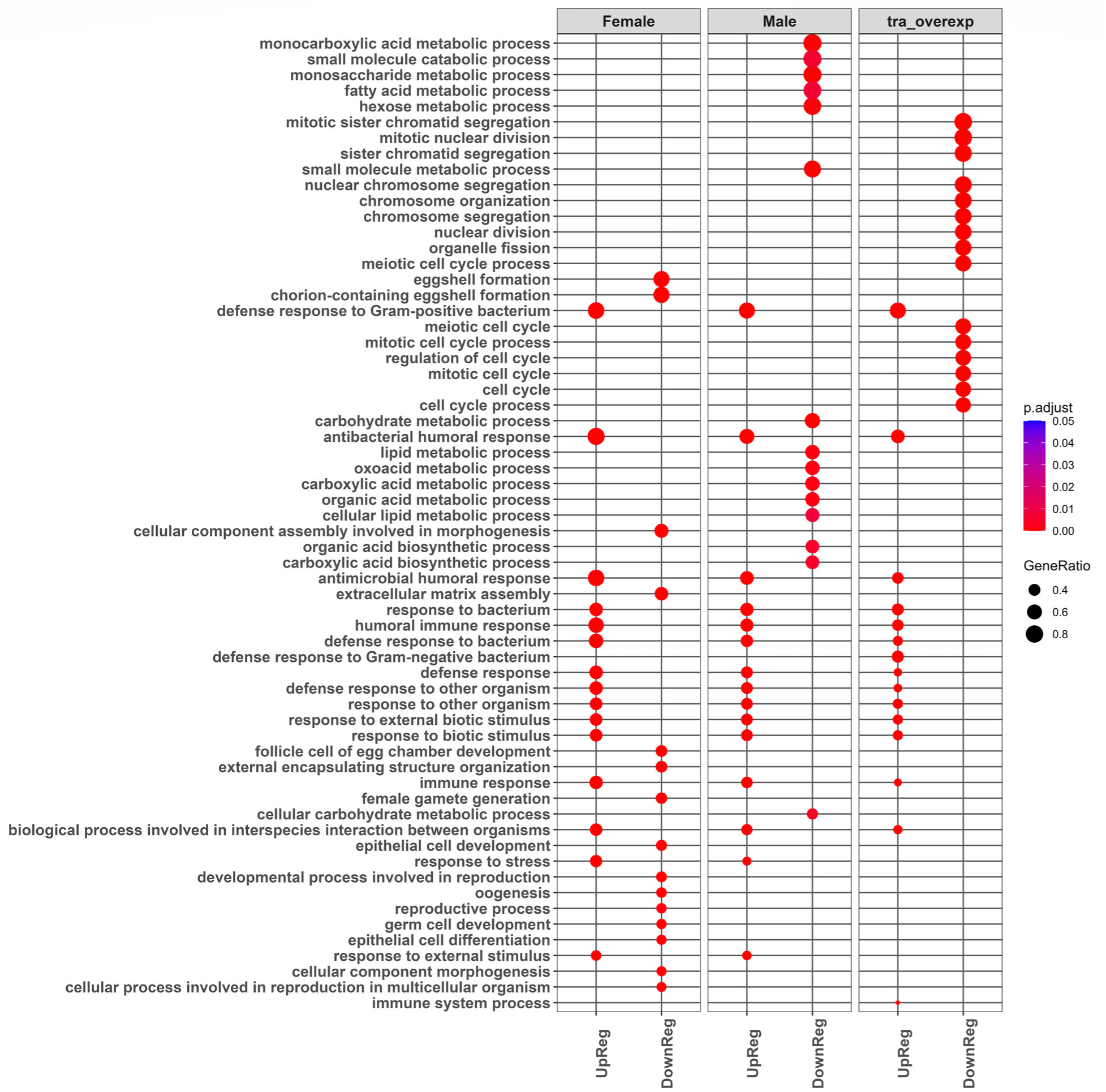
Top15 up and down-regulated Enriched Gene Ontology terms (gseGO: BP) in females, males, and *tra-*overexp upon *P. rettgeri* (PRET) infection (For details see. Supp. File 9). UpReg=Up Regulation, DownReg=Down Regulation.

### 3.7 Characterization of genes as regulated by or downstream of *tra*

The *tra-*mutant experimental analysis showed a substantial difference between females (XX) and *tra-*mutant (XX) animals with respect to both survival and DEGs. Moreover, the *tra-*mutant (XX), which is phenotypically similar to XY males in morphology and behavior, showed remarkable similarity with the males (XY) in the case of survival and DEGs. To investigate the role of *tra* in immune regulation, the interactions of treatment by sex and treatment by XX-genotype were used to identify the genes up and downstream of *tra* in the sex determination pathway (Table 1, Supp. File 7). This approach resulted in classifying 235 genes as upstream and 417 genes as regulated by or downstream of *tra*.

**Table 1:** **Genes sex differentially expressed** (For details, see Supp. File 7-8, Supp. Fig. 1) (Chang et al., 2011)

|  | *tra*-mutant | Total<br>Enriched GO: BP Terms | Immune-related<br>Enriched GO: BP Terms |
| --- | --- | --- | --- |
| <sup>1</sup>Upstream of *tra*<sup>F</sup> | 235 | 30 | 0 |
| <sup>2</sup>Downstream of *tra*<sup>F</sup> | 417 | 204 | 49 |
| <sup>3</sup>Sex-differentially expressed<sup>3</sup> | 652 |  |  |

1. FDR < 0.05 in female-male contrast and FDR > 0.05 in female-*tra*\_mutant contrast 2. FDR < 0.05 in female-male contrast and FDR < 0.05 in female-*tra*\_mutant contrast 3. FDR < 0.05 in female-male contrast

Enrichment analysis of the 235 upstream genes showed 30 enriched biological processes, which include no immune-related GO terms (Table 1, Supp. File 8). A large number of metabolic pathways and three terms related to cellular detoxification were also overrepresented in this group. Remarkably, the enrichment analysis of the 417 downstream genes showed 204 enriched biological processes, which include 49 immune-related GO terms. Further inquiry confirmed that most immune-related GO terms are linked to response against gram-negative bacterial infection (Supp. File 8). The enrichment analysis also identified genes of the Toll signaling pathway (GO:0008063) as being overrepresented among *tra-*regulated genes (Supp. File 8). The list of genes regulated by or downstream of *tra* also contained some genes of the core gram-negative response IMD pathway, but enrichment of the corresponding GO biological process term was insignificant.

### 3.8 Core systemic immune response pathways regulated by *tra*

Sex differences in the key ligand of the Toll pathways *spatzle* (*spz*) and principal membrane-bound toll receptor of the Tl gene family (Protein Toll) were classified as regulated by or downstream of the *tra* gene. The members of the *spatzle* activating pathway, *spirit, easter (ea),* and PGRP-SA, while showing sex differences in expression, but were classified as regulated upstream of *tra*. With respect to the regulation of the Toll pathway, several Serine protease inhibitors (Serpins) were also regulated by *tra*, including *Spn27A, Spn42Dd, and Spn88Ea*. It is worth mentioning that these specific Serpins also modulate another crucial defense response process, melanization, in *Drosophila*. Key Toll pathway members Pelle and Cactus were also identified as showing sex differences regulated downstream of *tra*. Expression of Toll-regulated effector molecules, Toll-regulated AMPs Drs (Drosomycin), and short-secreted peptides BomBc1 (Bomanins) were also classified as regulated by or downstream of *tra* (Lin et al., 2020). The Toll-induced hemolymph protein Bombardier (Bbd), which is required for bomanin function, was also sex differentially regulated under infection but appears to be regulated upstream of *tra* (Lin et al., 2020) (Supp. File 7).

Although genes in the IMD pathway were not overrepresented among tra-regulated genes, several key regulators were identified in our study as regulated by *tra*, which include the focal transcription factor of the IMD pathway NF-κB-Relish. This indicates the significant impact of *tra* regulation in the IMD pathway. In addition, *Charon,* which can directly activate the nuclear translocator NF-κB-Relish, showed sex differences regulated by or downstream of *tra*, enhancing the role of *tra* regulation. The analysis also found that Diap2/IAP2, an Inhibitor of apoptosis proteins and key mediator of NF-κB-Relish signaling, was also regulated by *tra*. The AMPs regulated downstream of the IMD pathway did not show distinct signatures of sex differential expression regulated by *tra*. Among them, only Drosocin (Dro) was identified in the downstream group, and Cecropin A1 (CecA1) was identified among genes with sex differences in immune gene expression regulated upstream of *tra* (Supp. File 7).

### 3.9 DEGs of IMD-Toll pathway in *tra*-mutant/overexpression

One of the core bacterial infection response pathways is the IMD Pathway in *Drosophila*. The pathway, including genes involved in signal reception, transduction, and cellular response, showed divergence among experimental groups of the *tra-*mutant and *tra-*overexpression in response to infection (Fig. 11-12, Supp. File 10). In the *tra-*mutant experiment, the analysis of the core systemic bacterial response pathways IMD and Toll genes showed all three groups, females, males, and *tra-*mutant flies, were upregulating these genes compared to wound control. However, the degree of upregulation diverges among these groups, which might determine the ultimate disease outcomes. In IMD pathways, the Attacin-AMPs were upregulated at similar levels for females, males, and *tra-mutants*, such as AttA, AttB, AttC, and AttD (Fig. 11, Supp. File 10). However, the Cecropin (Cec), Drosocin (Dro), and Diptericin (Dpt) AMP classes showed differences in the degree of upregulation (Fig. 11, Supp. File 10). Females (log2FC 6.7 to 4.5) upregulated the CecA1, CecA2, CecB, and CecC during infection, and the *tra-*mutant XX flies upregulated at a range of log2FC 5 to 3.9. Interestingly, the upregulation of these Cecropins (Cec) in the XX *tra-*mutant is more similar to the male (XY) (log2FC 5.6 to 3.7). Likewise, the upregulation of Drosocin (Dro) was higher in females (log2FC = 3.82) than in males and XX *tra-*mutant animals (*p*-adj = 0.014, 0.004). The upregulation of the males (log2FC=2.16) and XX *tra-*mutant animals (log2FC = 1.87) were similar for Drosocin (Dro). In the case of Diptericin (Dpt), females upregulated DptA (log2FC = 4.27) higher than *tra-*mutants (log2FC = 2.50) (*p*-adj = 0.015), and the male upregulated at log2FC = 3.24. DptB also showed upregulation in females (log2FC = 3.44), males (log2FC = 3.53), and *tra-*mutant (log2FC = 2.50). The central transcription factor of the IMD pathway, NF-kB-Relish expression, has the lowest level of upregulation in females (log2FC = 1.55) compared to males (log2FC = 2.33) and XX *tra-*mutant animals (log2FC = 2.93) which might indicate the initiation of immune suppression in females (*p*-adj = 0.016, 8.56E-05) (Fig. 11, Supp. File 10).

**Fig. 11:**
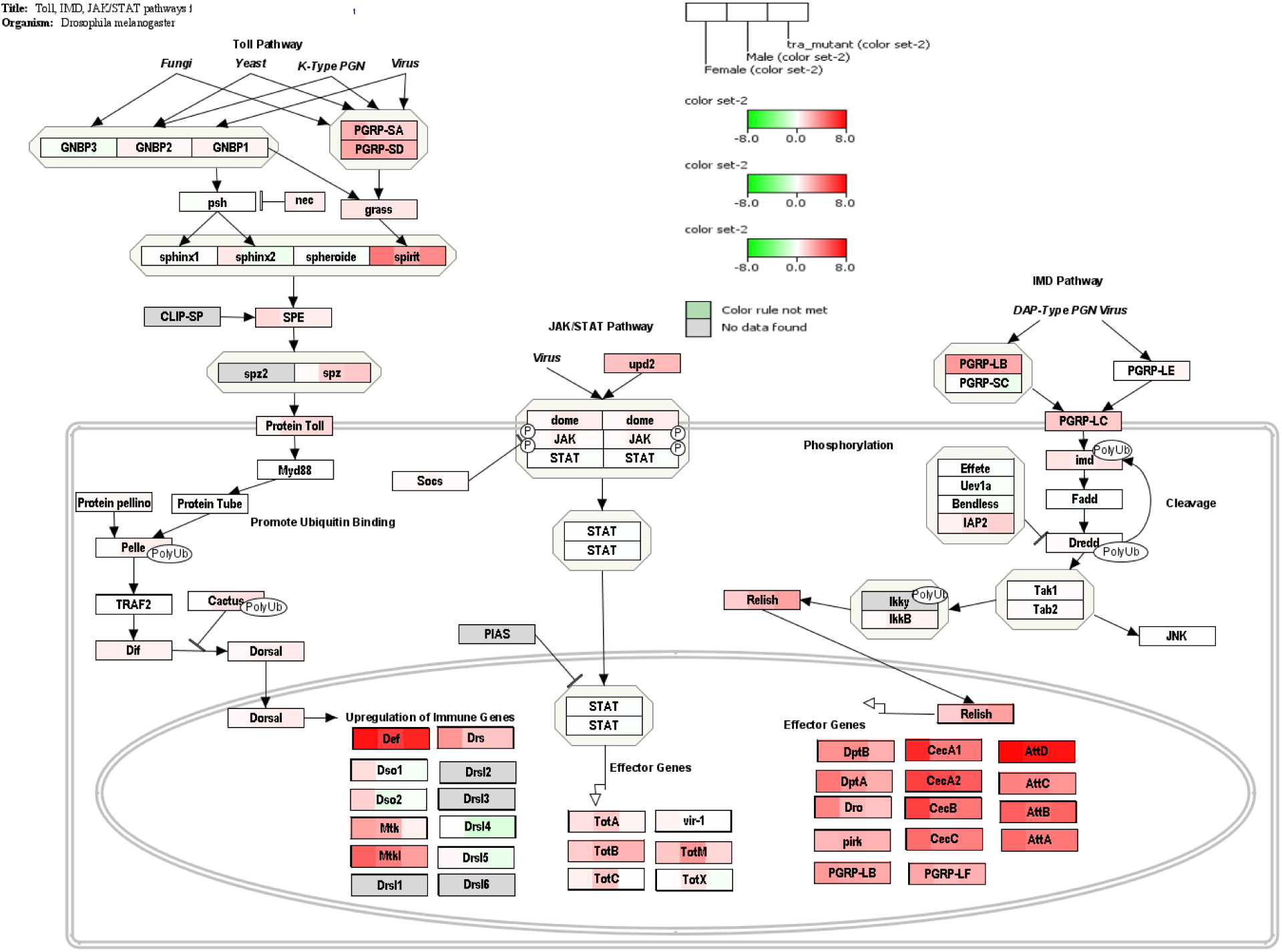
DEGs in females, males, and *tra-*mutant flies in response to *P. rettgeri* infection for genes in the Toll-IMD-Jak/Stat signaling pathway (wiki pathway ID: WP3830). The expression (log2FoldChange) of the pathway is represented without statistical cut-off. Red represents upregulation, green represents downregulation, and white represents no change compared to wound control (For details, see Supp. File 10, Supp. Fig. 2).

The Toll pathway also had distinct patterns of expression in females compared to Tra^F,^ lacking males and XX *tra-*mutants. The core ligand *spatzle* (*spz*) and Spz binding receptor Toll (*Tl*) were significantly upregulated in males (*p*-adj = 1.19E-06, 0.01) and *tra-*mutant (*p*-adj = 3.03E-08, 0.003) compared to females. Females did not upregulate primary transcription factor NF-kB-Dorsal (*dl*) of the Toll pathway compared to wound control, whereas both in males (log2FC = 0.50) and *tra-*mutant (log2FC = 0.61) upregulated. However, *spz* activating pathways elements such as Spatzle-Processing Enzyme (SPE) and Spirit were upregulated by females (log2FC = 1.13; 4.22), males (log2FC = 0.61; 2.67) and *tra-*mutants (log2FC = 0.56; 3.65). In the case of the AMPs of the Toll pathway, Def, Drs, and Mtkl were upregulated by female (log2FC=3.2 to 7.4), male (log2FC=1.77 to 6.1), and *tra-*mutants (log2FC=1.81 to 6.82) (Fig. 11, Supp. File 10).

In the *tra-*overexpression experiment, the analysis of core systemic bacterial response pathways IMD and Toll revealed that the level of upregulation differs among each of the two sexes relative to the XY *tra-*overexpression animals (Fig. 12, Supp. File 10). The data of the *tra-*overexpression study showed that *tra-*overexpressing animals significantly upregulated the genes of the IMD pathways compared to both the females and males, including many of the AMPs. The interaction analysis comparing changes in expression under infection among females, males, and XY *tra-*overexpression animals identified the AMPs of the IMD pathway DptA, DptB, and AttC were significantly upregulated in *tra-*overexpression animals than both sexes (*p*-adj < 0.05) (Fig. 12, Supp. File 10). *tra*-overexpression flies also significantly upregulated the CecB than males (*p*-adj < 0.05). Further, *tra*-overexpressed animals showed a significant upregulation of the AttA and AttB compared to females (*p*-adj < 0.05). Furthermore, tra-overexpression led to the upregulation of CecA1 (log2FC = 5.60), CecC (log2FC = 4.87), and Dro (log2FC = 3.39), with females exhibited upregulation in CecA1 (log2FC = 5.01), CecC (log2FC = 4.30), and Dro (log2FC = 2.91). *tra-overexpressed* flies also significantly upregulated key genes of IMD such as transcription factor NF-kB-Relish, regulator IAP2, membrane-bound core signal receptor PGRP-LC, and ex*tra*cellular PGRP-LB and the interaction showed that these upregulations were significantly higher from that in females (*p*-adj <0.05) (Fig. 12, Supp. File 10).

**Fig. 12:**
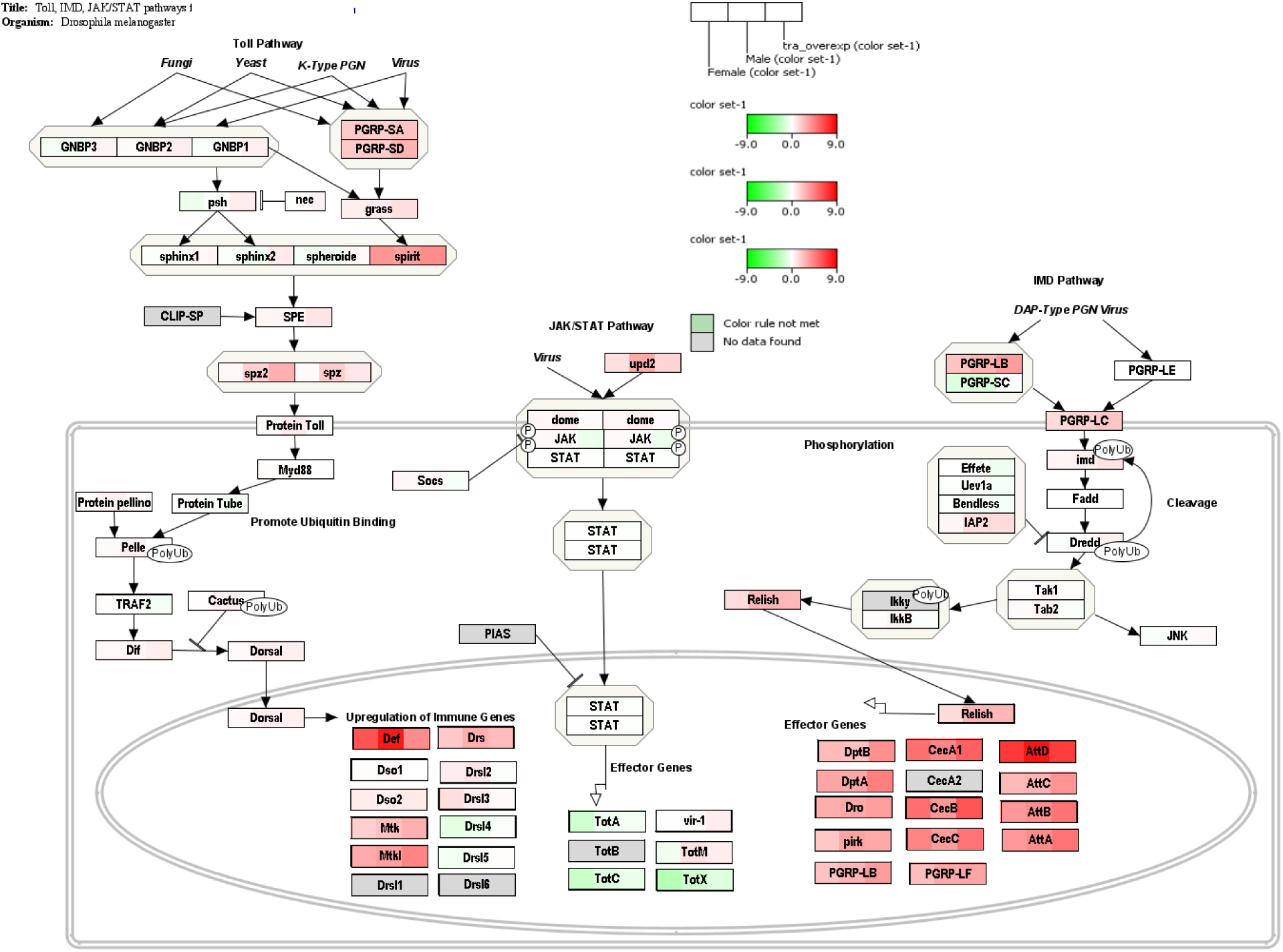
DEGs in females, males, and *tra-*overexpressed flies in response to *P. rettgeri* infection for genes in the Toll-IMD-Jak/Stat signaling pathway (wiki pathway ID: WP3830). The expression (log2FoldChange) of the pathway is represented without statistical cut-off. Red represents upregulation, green represents downregulation, and white represents no change compared to wound control (For details, see Supp. File 10, Supp. Fig. 2).

In *tra*-overexpression, the contrast between females and males showed the two sexes upregulated the AMPs and other IMD genes similarly (only IAP2 was significantly upregulated in females), which might indicate that females controlled the infection at an earlier time point and, therefore, initiated IMD pathway suppression which contributes to the absence of sex differences between these sexes (Fig. 12, Supp. File 10). This pattern is also found in other studies of immune challenges (Khan2023_Unpublished_work_1, 2023; Khan2023_Unpublished_work_2, 2023). Upregulation of the key transcription factor NF-kB-Relish in males was significantly higher than in females (*p*-adj <0.05); this might suggest males are still inducing IMD pathway-regulated genes to control the bacterial infection at 16hpi, an occurrence that is consistent with the survival and bacterial load. This pattern of males and females was also visible in our previous study of bacterial infection (Khan2023_Unpublished_work_1, 2023; Khan2023_Unpublished_work_2, 2023) (Fig. 12, Supp. File 10).

In the *tra*-overexpression experiment, Toll pathway genes were similar in comparisons between females and the XY *tra*-overexpression individuals. *tra-*overexpression animals upregulated only Mtk at a higher level relative to females (*p*-adj < 0.05) (Fig. 12, Supp. File 10). Similarly, females elevated only one AMP of Toll (Def) as compared to *tra-*overexpression animals (*p*-adj < 0.05). *tra-*overexpression resulted upregulation of Mtkl (log2FC = 4.28), PGRP-SA (log2FC = 2.10), PGRP-SD (log2FC = 2.63) and spirit (log2FC = 4.02) and while females exhibited upregulation in Mtkl (log2FC = 2.94), PGRP-SA (log2FC = 1.84), PGRP-SD (log2FC = 1.86) and spirit (log2FC = 3.68).

Def had the highest upregulation of any genes in the *tra-*overexpression study. When compared to the wound control, males upregulated Def 16-Fold, whereas XY *tra-*overexpression animals and females elevated expression of this gene ∼8Fold and ∼12Fold, respectively (Fig. 12, Supp. File 10). In addition, remarkably, the male showed significantly higher upregulation of the core ligand of the Toll signaling *spz* and Toll-AMPs like Drs and Def relative to both females and *tra-*overexpression groups (*p*-adj <0.05), which again supports the assumptions that the males were actively controlling the infection whereas *tra*-overexpressed and females might have initiated suppression and passed the highly activated phase of the immune response.

## 4. Discussion

A limited number of studies have been conducted on complex sexually dimorphic traits such as immune response, mainly due to the inherent variability and complexity of the immune response itself (Lazzaro & Little, 2009). In this study, we aimed to evaluate the role of key sex determination pathway gene *tra* in immune sex differences by examining XX *tra-*mutant and XY *tra-*overexpression animals in *Drosophila*. The absence of functional Tra^F^ protein in XX flies results in significantly lower survival relative to typical XX female flies. The overexpression of Tra^F^ confirmed the role of *tra* in enhancing the survival of bacterial infections. When *tra* was ubiquitously overexpressed, XY individuals exhibited notably higher survival rates, which were, in fact, higher than both XX females and XY males. Genes with sex differences in expression due to bacterial infection regulated by or downstream of *tra* were also identified. These genes revealed that *tra* plays a regulatory role in various immunological processes, including the canonical bacterial infection response pathways of Drosophila, Toll, and IMD signaling pathway genes. In addition, the analysis of differentially expressed genes (DEGs) in *tra-*mutant XX individuals confirmed their inability to mount an effective immune response compared to individuals with the natural presence of Tra^F^. On the other hand, *tra-*overexpression XY animals had a surprisingly heightened ability to mount an immune response in comparison to both males and females, revealing that this may be an excellent model for understanding the role of *tra* in regulating the immune response.

### 4.1 Tra regulates immune differences between males and females and promotes survival

The survival of an organism following infection is often interpreted as a direct measure of the effectiveness of the immune response. The survival assay revealed that the absence of Tra^F^ in XX flies results in a significant decrease in survival rate. The rate of survival was detected to be the same as that observed in males (Fig. 3). To further confirm the role of Tra^F^ in the regulation of the immune response, *tra* was overexpressed in XY animals, where Tra^F^ is not naturally produced. The results showed significantly higher survival of the XY *tra-*overexpression individuals, and it was higher than in females where Tra^F^ is expressed in the natural pattern (Fig. 3). This study is the first report of the immune response in *tra-*mutant and *tra-*overexpression animals.

Tra is required for sex differences that are regulated by the transcription factors Fru and Dsx. It is known to regulate sex dimorphism in complex traits. For example, *tra* regulates sex differences in body size in Drosophila, with *tra-*mutant XX flies having a significant reduction in pupal volume compared to wild-type female flies and in comparable levels to wild-type males, which is 30% smaller than females in nature (Rideout et al., 2015). In addition, the same study also performed ubiquitous overexpression of *tra* using the Daughterless-GAL4 driver in XY animals. They found that pupal volume exceeded the size of the wild-type male, and *tra-*overexpression flies were significantly larger than wild-type females (Rideout et al., 2015). Our findings of differences in survival rate in both *tra-*mutant and *tra-*overexpressed flies compared to individuals sharing the same sex chromosome genotype showed a similar pattern, with XX *tra*-mutant animals being most similar to males and XY *tra*-overexpression animals having greater survival than both XX females and XY males. Interestingly, in both cases, the effect of the ubiquitous overexpression is a surplus to the natural level of the trait. This may reflect the mechanism by which *tra* regulates sex differences in complex traits.

The bacterial load in *tra-*overexpression XY flies was significantly lower than XY males at 16hpi; however, the XX *tra-*mutant load was not significantly different from XX females. Thus, the bacterial load data is partly consistent with survival; this might happen because the infection is still progressing at 16hpi, and taking a snapshot of the whole process reveals only part of the effect of the *tra-*mutation. In addition, the bacterial load differed between the sexes, alongside variation in the individual capacity to control the infection (Duneau, Ferdy, et al., 2017).

### 4.2 Immune gene expression in *tra*-mutant and *tra*-overexpressing animals

The differential expression profile of *tra*-mutant flies confirmed the essential role of *tra* in sex differences in immunity. The *tra-*mutant XX animals were unable to mount immune gene expression at the level of wild-type females; instead, it is more comparable to the wild-type males, which also lack expression of Tra^F^. At 16hpi, females started to control activation of the IMD and Toll pathways, whereas both males and *tra-*mutants were actively attempting to upregulate effectors regulated by Toll and IMD. Animals in which Tra^F^ is absent, male and *tra-*mutant, upregulated key elements of IMD and Toll to increase the effectiveness of the immune response. In the IMD pathway, the males and *tra-*mutant animals upregulated the central transcription factor of the IMD pathway NF-kB-Relish around 5- and 6-Fold, respectively, compared to ∼3Fold in females, which might reflect the initiation of negative regulation of these pathways in the females. The higher upregulation of Relish during active immune response was observed in previous studies (Khan2023_Unpublished_work_1, 2023; Vincent & Dionne, 2021). An identical pattern was observed in the expression of the three central components of the Toll pathways: core ligand *spatzle* (*spz*), Spz binding receptor Toll (*Tl*), and primary transcription factor NF-kB-Dorsal (*dl*). While males and *tra*-mutants significantly upregulated *spatzle* and Toll (Tl) genes, females did not exhibit significant upregulation of these genes. This is consistent with the upregulation of key components of the Toll pathway during the immune response to infection (Duneau, Kondolf, et al., 2017; Khan2023_Unpublished_work_1, 2023; Khan2023_Unpublished_work_2, 2023). Although females may have initiated downregulation of IMD and Toll, the number of AMPs such as Drosocin (Dro) and Diptericin (DptA) maintained increased expression levels in females, whereas *tra-*mutant flies expressed these genes at lower levels than females even in the active state of infection control. Likewise, in AMPs of the Toll pathway, females upregulated Drs greater than males and XX *tra-*mutants.

Overall, the results suggest that at 16hpi, females initiate suppression of immune pathways, such as IMD and Toll, due to the higher capacity of infection control. The survival analysis data supports the assumption that the presence of *tra* provides females with an advantage in efficiently controlling bacterial infections compared to both males and *tra-*mutant flies, which lack Tra^F^. Our experimental data indicate that the highly activated state of upregulation of effectors in females was earlier than 16hpi. This points to the need for studies that capture these responses over multiple time points, which may reveal the dynamics of the immune response and the full complexity of the role of *tra* in regulating sex differences in immunity.

The role of *tra* as a central regulator of immune sex dimorphism is also supported by the *tra-*overexpression experiment. Animals that express Tra^F^ upregulate effectors of the IMD pathway at levels much higher than the males; however, the lower level of upregulation of Relish indicates the initiation of IMD pathway suppression in both females and *tra-*overexpression flies, whereas the IMD response in males remains activated (Khan2023_Unpublished_work_1, 2023; Khan2023_Unpublished_work_2, 2023). Overall, the contrast of the male with females and *tra-*overexpression animals, which both express Tra^F^ protein, suggests that *tra-*overexpression flies may control the infection at an earlier time point relative to females and males. *Tra-overexpression* flies evidently suppressed the IMD pathway via the upregulation of negative regulators such as PGRP-LF (log2FC=3.14) and *pirk* (log2FC=3.01); still *tra-*overexpressing flies significantly differed from males who were actively upregulating IMD pathways at 16hpi (male, Relish:log2FC = 2.57); whereas none of the IMD-genes in females were significantly upregulated relative to males at 16hpi (except IAP2) (Fig. 10, Supp. File 10). Again, this assumption of high IMD pathway activation in *tra-*overexpression animals is supported by the unusually high upregulation of *Relish* (log2FC = 2.57) (equal to male log2FC = 2.57, 2x higher than female log2FC = 1.22) at 16hpi even though the *tra-*overexpression flies were controlling the infection efficiently according to survival data and bacterial load (Fig. 10, Supp. File 10).

### 4.3 Tra regulates sex differences in the Toll and IMD pathways

The regulatory targets of *tra* were identified by comparing differential gene expressions between sexes and *tra-mutants*. Genes with sex differences regulated upstream of *tra* were enriched for mostly metabolic processes, while genes regulated by or downstream of tra were enriched for bacterial infection-related processes. Interestingly, the enriched biological process list contains the core bacterial response pathway Toll signaling, which provides support to the previous report of Toll-mediated regulation of the immune sex dimorphism (Duneau, Kondolf, et al., 2017).

Although IMD, another canonical bacterial defense response pathway, was not identified among the enrichment terms, sex differences in the expression of key genes such as transcription factor NF-kB-Relish are regulated by *tra*. Interestingly, the results also underscore the importance of the IMD pathway as one of the key regulators of immune sex dimorphism in Drosophila (Vincent & Dionne, 2021).

Although our findings identified both Toll and IMD pathway components as regulated by or downstream of *tra*, several negative regulators of Toll signaling pathways, such as Cactus, Spn42Dd, and Spn88Ea, are also regulated by *tra*. Surprisingly, several Toll-related genes, which are regulated downstream of the *tra*, are connected to the fungal response. For instance, the only AMP, Drosomycin (Drs), found to be directly regulated by *tra*, is specialized in the response to fungal infection, and both Spn42Dd and Spn88Ea function as negative regulators during fungal attack.

In summary, *tra* has a regulatory role that effects the expression of the Toll pathway, Toll-related Bomanins, IMD, and melanization cascade, suggesting that there is likely not a single biological pathway or process that explains the complexities of sex differences in immunity.

### 4.5 Functions impacted under infection in *tra*-mutant and *tra*-overexpression animals

The gene set enrichment analysis of DEGs (FDR 0.05 and 2FC) showed the *tra-*mutant consistently downregulated genes involved in mitochondrion organization, whereas females and males downregulated several other processes during infection. The downregulation of the metabolic and reproductive process upon infection has been observed in our prior study (Khan2023_Unpublished_work_1, 2023). The *tra-*mutant upregulated genes involved in immunological processes robustly according to the enrichment analysis, which might indicate a higher stress level due to bacterial infection relative to females and males. In addition, the constant overstimulation of the immune system might impact the overall physiological events and survival of an animal (Hoffmann, 2003; Tanji & Ip, 2005). Our previous studies observed that flies that controlled bacterial infection upregulated many genes with functions related to negative regulation of the immune response, putatively to reduce damage resulting from overstimulation (Khan2023_Unpublished_work_1, 2023; Khan2023_Unpublished_work_2, 2023). Downregulation of genes involved in metabolic processes and upregulation of genes involved in negative regulation of the immune response was completely absent in the *tra-*mutant. The opposite scenario was observed in the *tra-*overexpression animals, where the GO analysis of DEGs (FDR 0.05 and 2FC) showed a remarkable 333 terms out of 399 enriched terms corresponding to downregulated genes. Among these terms were many related to reproduction, development, and cell division, which might suggest efficient strategies to handle bacterial infection considering energy trade-offs between reproduction and immunity (Metcalf et al., 2020; Schwenke et al., 2016). Furthermore, the upregulation of several negative regulatory processes impacting the immune response strengthens our previous assumption regarding the initiation of bacterial response suppression in *tra-*overexpression individuals at 16hpi.

## 5. Conclusion

The absence of a sex determination pathway gene *transformer* results in reduced effectiveness of the immune response to gram-negative extracellular bacterial infection. The differential gene expression analysis confirmed lower levels of upregulation of effectors, such as AMPs, in the *tra*-mutant animals. Moreover, ubiquitously overexpressed *tra* resulted in much higher survival relative to both wild-type females and males. The transcriptomic profiles also indicate higher efficacy of the *tra-*overexpressed flies upon bacterial challenge. The downstream targets of *tra* showed significant enrichment of genes involved in immune processes, with a significant impact of *tra* on both Toll and IMD signaling pathways. Overall, this study found a significant impact of *tra* on sex dimorphism of the immune response and on immune response as a whole; however, whether *tra* directly or indirectly regulates these genes remains an open question. Tra regulates the key transcription factors *doublesex* and *fruitless* in Drosophila; it will be important to determine whether *doublesex* or *fruitless* has a role in regulating sex differences in the immune response. This will illuminate our understanding of whether *tra* uses the *sxl-tra-dsx^F^* cascade to regulate the immune sex dimorphism or some other regulatory cascade.

## Supporting information

Supplement_File_1_QC_Report_Novagen

Supplement_File_2_tra_STAR_Mapping_Stat

Supplement_File_3_globalDEresultsTra_full_model#mutant17869#overexp17869

Supplement_File_4_globalDEresultsTra_full_model#mutant13694#overexp13596

Supplement_File_5_globalDEresultsTra_full_model_with_symbols#mutant13649#overexp13553

Supplement_File_6_Tra_full_model_#mutant_#overexpression_counts

Supplement_File_7_Tra_mutant_upstream#235_downstream#417_gene_list

Supplement_File_8_Tra_mutant_upstream#235_downstream#417_enrichGO#upstream30#downstream#204

Supplement_File_9_gseGO_#mutant13694_overexp13596_pretVSpbs_FDR0.05_2FC

Supplement_File_10_All_Functional_Gene_Class#mutant13649#overexp13553

## Supplement Figure and Tables

**Supp. Fig. 1:**
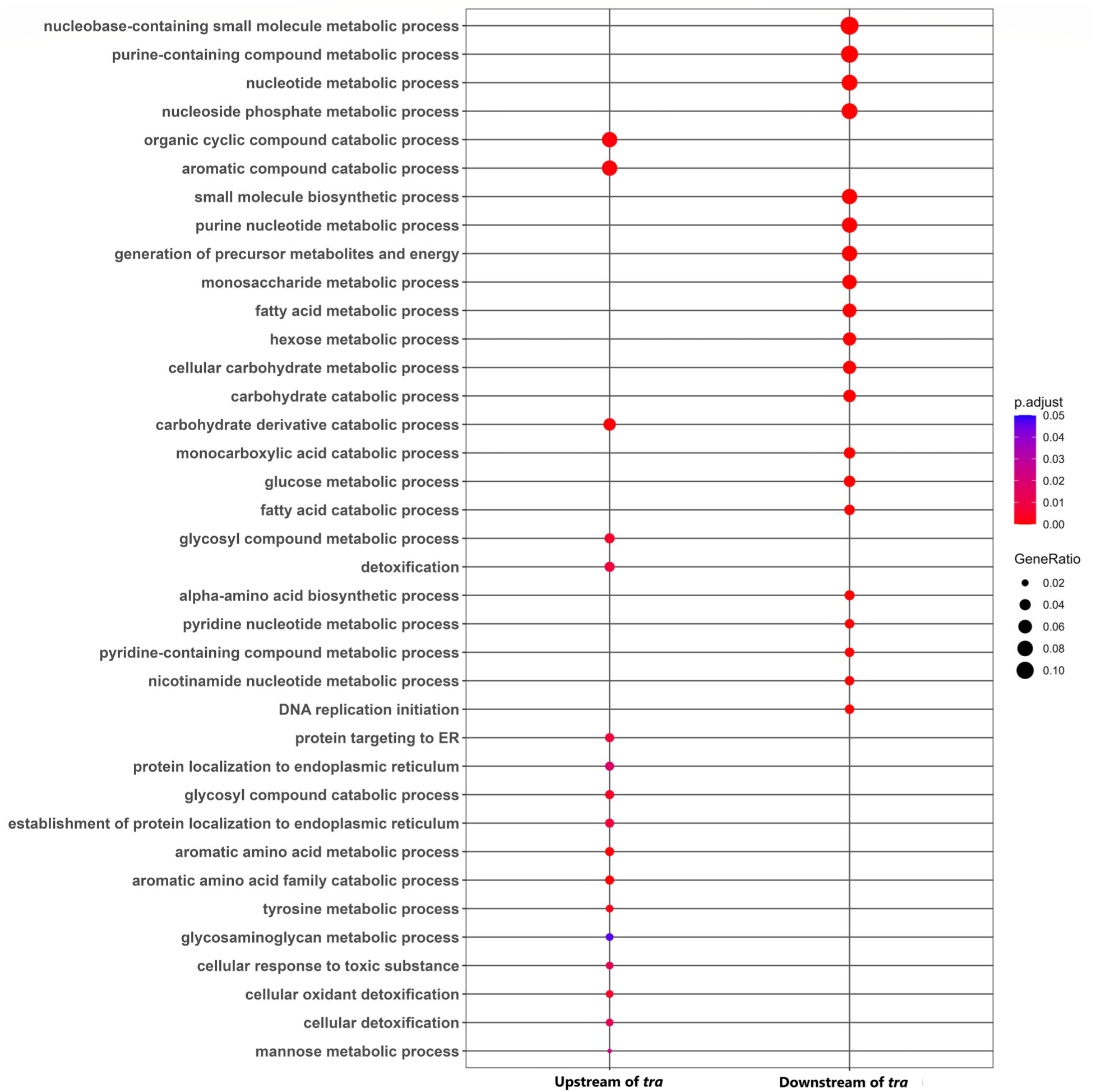
Top20 (Unique) Enriched Gene Ontology terms (enrichGO: BP) in upstream (17) and downstream (20) of *tra* upon *P. rettgeri* (PRET) infection based on *tra-*mutant experiment. (Note: 17 terms for the upstream of *tra*) (For details, see Supp. File 7-8).

**Supp. Fig. 2:**
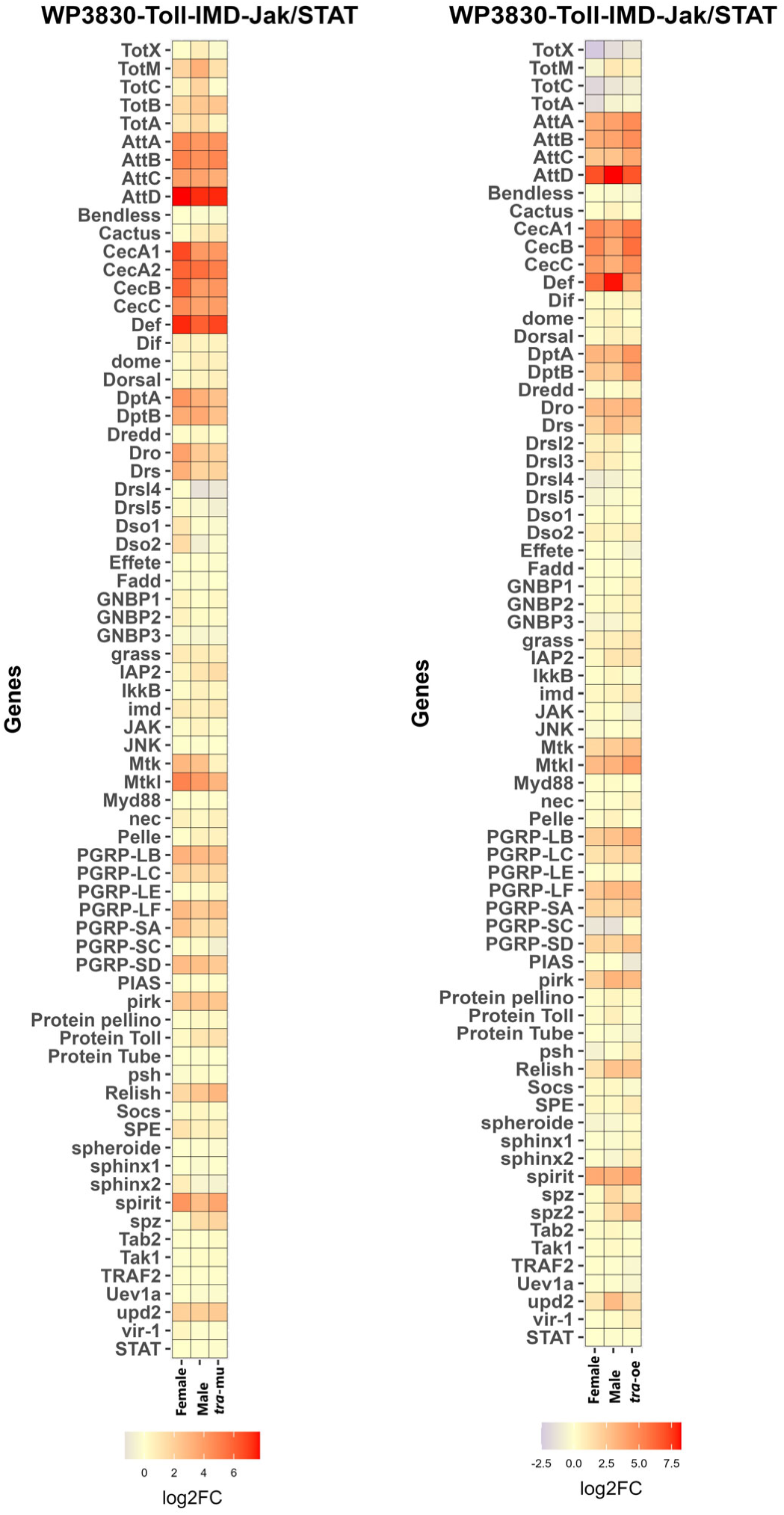
Sex differences in the Toll-IMD-Jak/Stat signaling pathway (wiki pathway ID: WP3830) in *tra-*mutant and *tra-*overexpression experimental flies. The expression (log2FC) of the pathway is represented without statistical cut-off. Female = wild type Female, Male = wild type Male, *tra-*mu=*tra-*mutant, *tra-*oe = *tra-*overexpressed. The expression (log2FC) of the pathway is represented without statistical cut-off (For details, see Supp. File 10).

