## Supplement_File_1_QC_Report_Novagen for "Regulation of sex differences in innate immunity by sex-determining gene *transformer* in *Drosophila melanogaster*"

A. Library Preparation and Sequencing
1 Sample Quality Control
2 Library Construction, Quality Control and Sequencing
B. Results and Instructions
1 Data Quality Control
1.1 Distribution of Sequencing Quality
1.2 Distribution of Sequencing Error Rate
1.3 Distribution of A/T/G/C Base
1.4 Results of Raw Data Filtering
2 Summary of Sequencing Data Information
C. Appendix
1 Introduction of Sequencing Data Format
2 Explanation of Sequencing Data Related
3 References

### QC Analysis Report

| Contract Information | Contract Content |
| --- | --- |
| Contract ID | H202SC22092632 |
| Contract Name | Davis-US-Auburn-48-Drosophila melanogaster-EukmRNAseq-6G-WOBI-NVUS2022090740 |
| Batch ID | X202SC22092632-Z01-F001 |
| Report Time | 2023-02-21 |

Novogene Co., Ltd

### A. Library Preparation and Sequencing

From the RNA samples to the final data, each step, including sample test, library preparation, and sequencing, influences the quality of the data, and data quality directly impacts the analysis results. To guarantee the reliability of the data, quality control (QC) is performed at each step of the procedure. The workflow is as follows:

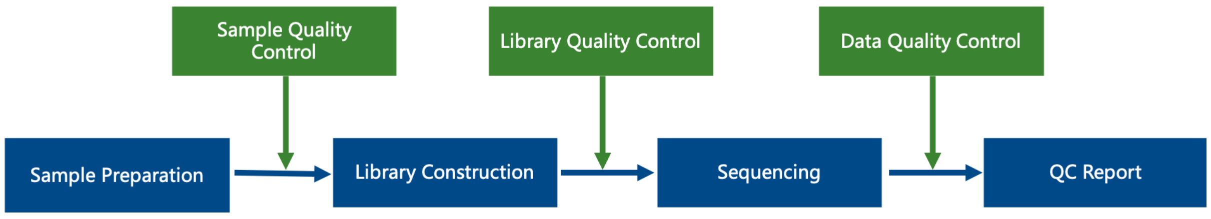


#### 1 Sample Quality Control

Please refer to Novogene's QC report for methods of sample quality control.

#### 2 Library Construction, Quality Control and Sequencing

Messenger RNA was purified from total RNA using poly-T oligo-attached magnetic beads. After fragmentation, the first strand cDNA was synthesized using random hexamer primers followed by the second strand cDNA synthesis. The library was ready after end repair, A-tailing, adapter ligation, size selection, amplification, and purification. Following is the workflow of library construction:

A. Library Preparation and Sequencing
1 Sample Quality Control
2 Library Construction, Quality Control and Sequencing
B. Results and Instructions
1 Data Quality Control
1.1 Distribution of Sequencing Quality
1.2 Distribution of Sequencing Error Rate
1.3 Distribution of A/T/G/C Base
1.4 Results of Raw Data Filtering
2 Summary of Sequencing Data Information
C. Appendix
1 Introduction of Sequencing Data Format
2 Explanation of Sequencing Data Related
3 References

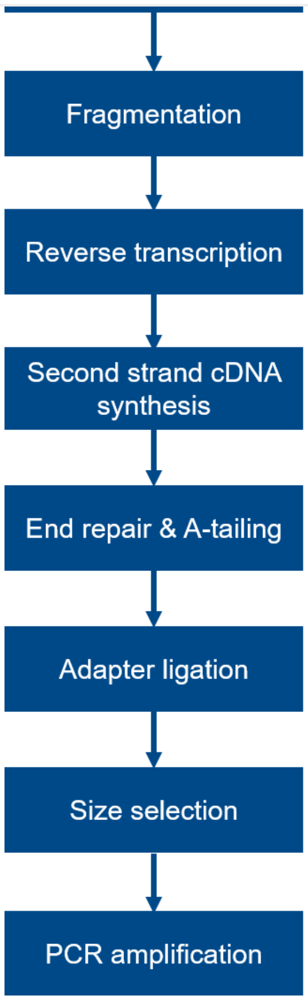


The library was checked with Qubit and real-time PCR for quantification and bioanalyzer for size distribution detection. Quantified libraries will be pooled and sequenced on Illumina platforms, according to effective library concentration and data amount.

Novogene Co., Ltd

### B. Results and Instructions

#### 1 Data Quality Control

##### 1.1 Distribution of Sequencing Quality

The “e” represents the sequence error rate and $Q_{phred}$ represents the base quality value,$Q_{phred}=-10\log_{10}(e)$. The relationship between sequencing error rate (e) and sequencing base quality value ($Q_{phred}$) is shown in the table below:

| Phred score | Error base | Right base | Q-score |
|---|---|---|---|
| 10 | 1/10 | 90% | Q10 |
| 20 | 1/100 | 99% | Q20 |
| 30 | 1/1000 | 99.9% | Q30 |
| 40 | 1/10000 | 99.99% | Q40 |

The distribution of quality score is shown in **Fig.1**:

A. Library Preparation and Sequencing
1 Sample Quality Control
2 Library Construction, Quality Control and Sequencing
B. Results and Instructions
1 Data Quality Control
1.1 Distribution of Sequencing Quality
1.2 Distribution of Sequencing Error Rate
1.3 Distribution of A/T/G/C Base
1.4 Results of Raw Data Filtering
2 Summary of Sequencing Data Information
C. Appendix
1 Introduction of Sequencing Data Format
2 Explanation of Sequencing Data Related
3 References

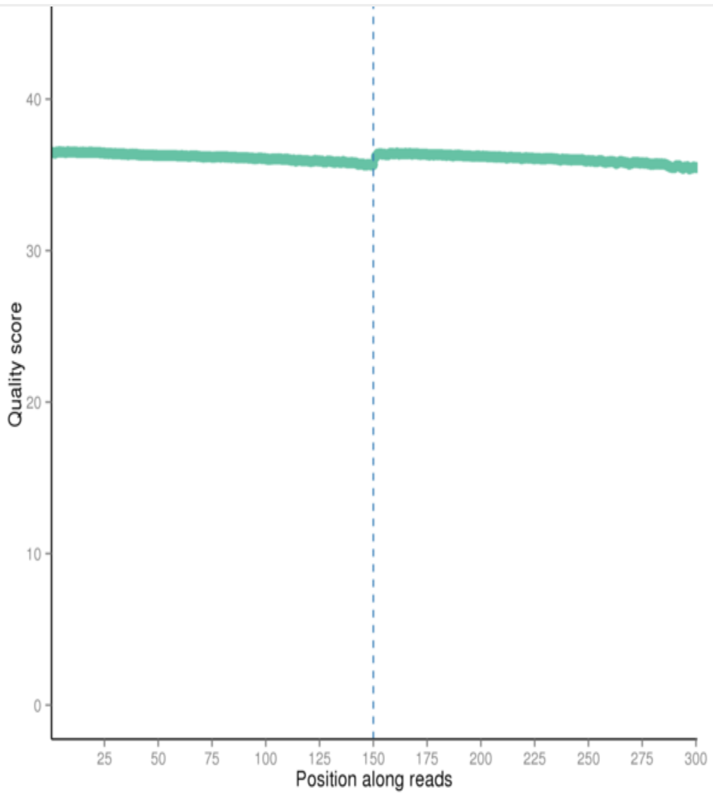


T3
T4
T8
T11
T17
T20
T24
T32
T33
T35

**Fig.1 Distribution of Sequencing Quality**

The base position is on the horizontal axis and the sequencing quality is on the vertical axis.

Novogene

Novogene Co., Ltd

#### 1.2 Distribution of Sequencing Error Rate

For Illumina SBS technology, the distribution of sequencing error rate has two features:

(1) Error rate grows with sequenced reads extension because of the consumption of sequencing reagent. The phenomenon is common in the Illumina high-throughput sequencing platform (Erlich Y. et al. 2008; Jiang et al. 2011).

(2) The reason for the high error rate of the first six bases is that the random hex-primers and RNA template bind incompletely in the process of cDNA synthesis (Jiang et al.2011).

The error rate of this project is shown in **Fig.2**:

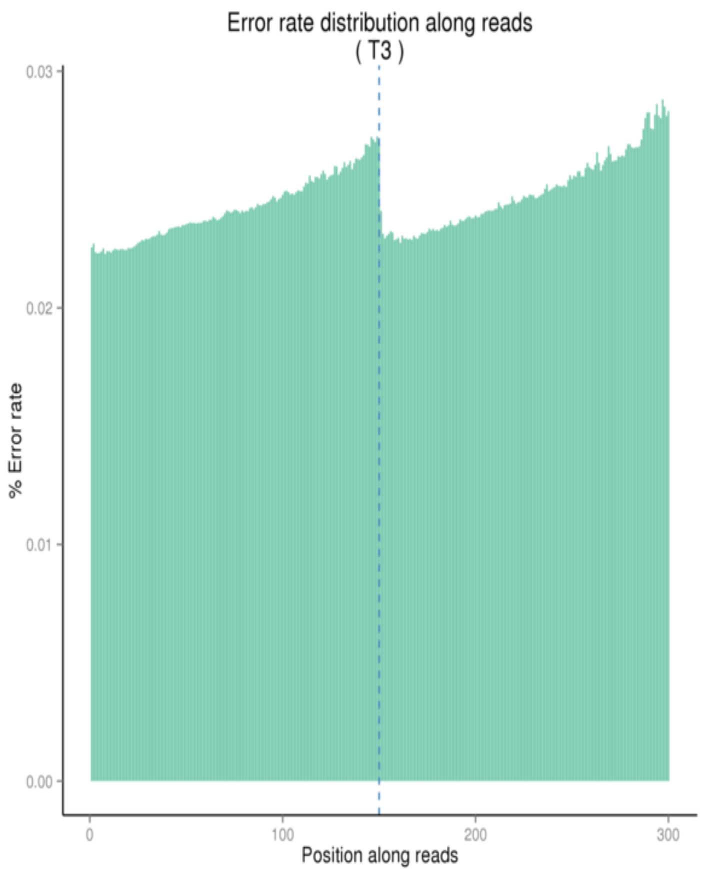


Search..
T3
T4
T8
T11
T17
T20
T24
T32
T33
T35

The base position is on the horizontal axis and the single base error rate is on the vertical axis


A. Library Preparation and Sequencing
1 Sample Quality Control
2 Library Construction, Quality Control and Sequencing
B. Results and Instructions
1 Data Quality Control
1.1 Distribution of Sequencing Quality
1.2 Distribution of Sequencing Error Rate
1.3 Distribution of A/T/G/C Base
1.4 Results of Raw Data Filtering
2 Summary of Sequencing Data Information
C. Appendix
1 Introduction of Sequencing Data Format
2 Explanation of Sequencing Data Related
3 References




#### 1.3 Distribution of A/T/G/C Base

It is used to identify the separation situation of AT and GC by checking the distribution of GC content. According to the principle of complementary bases, the content of AT and GC should be equal at each sequencing cycle and be constant and stable in the whole sequencing procedure. But in practical measurement, due to the primer amplification bias and some other reasons, the first 6 to 7 nucleotides will fluctuate which is normal and reasonable.

The distribution of GC content is shown in **Fig.3**:

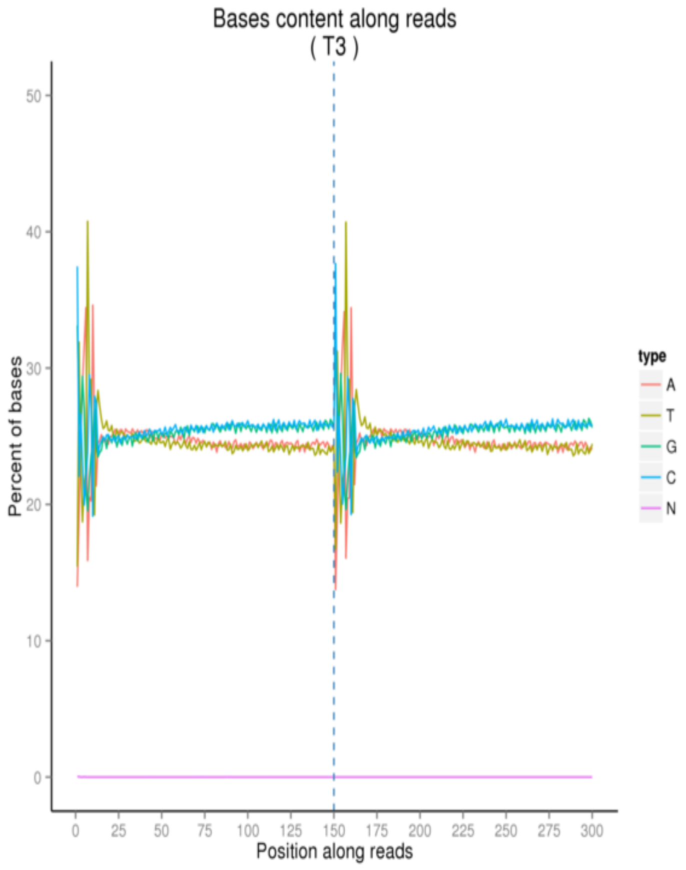


Search..

T3
T4
T8
T11
T17
T20
T24
T32
T33
T35

**Fig.3 A/T/G/C Distribution**

The base position is on the horizontal axis and the single base percentage is on the vertical axis



#### 1.4 Results of Raw Data Filtering

The sequenced reads (raw reads) often contain low quality reads and adapters, which will affect the analysis quality. So it's necessary to filter the raw reads and get the clean reads. The filtering process is as follows:

(1) Remove reads containing adapters.

(2) Remove reads containing N > 10% (N represents the base cannot be determined).

(3) Remove reads containing low quality (Qscore<= 5) base which is over 50% of the total base.

**Sequences of adapter**

5' Adapter:

5'-AGATCGGAAGAGCGTCGTGTAGGGAAAGAGTGTAGATCTCGGTGGTCGCCGTATCATT-3'

5'-GATCGGAAGAGCACACGTCTGAACTCCAGTCACGGATGACTATCTCGTATGCCGTCTTCTGCTTG-3'


A. Library Preparation and Sequencing
1 Sample Quality Control
2 Library Construction, Quality Control and Sequencing
B. Results and Instructions
1 Data Quality Control
1.1 Distribution of Sequencing Quality
1.2 Distribution of Sequencing Error Rate
1.3 Distribution of A/T/G/C Base
1.4 Results of Raw Data Filtering
2 Summary of Sequencing Data Information
C. Appendix
1 Introduction of Sequencing Data Format
2 Explanation of Sequencing Data Related
3 References


The Sequencing data filtration of this project can be seen in **Fig.4** :

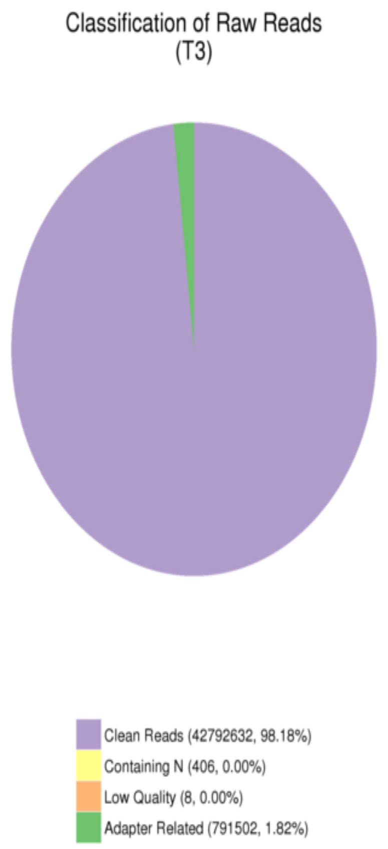



Search..
T3
T4
T8
T11
T17
T20
T24
T32
T33
T35


**Fig.4 Composition of Raw Data**

Different color for different components:

(1)Adapter related: (reads containing adapter) / (total raw reads)

(2)Containing N: (reads with more than 10% N) / (total raw reads)

(3)Low quality: (reads of low quality) / (total raw reads)

(4)Clean reads: (clean reads) / (total raw reads)



#### 2 Summary of Sequencing Data Information

The total output of data on the sequencer: Raw data 556.2 G.

The detail statistics for the quality of sequencing data are shown in **Table 1**.

**Table 1 Data Quality Summary**

| Sample | Raw reads | Raw data | Effective(%) | Error(%) | Q20(%) | Q30(%) | GC(%) |
|---|---|---|---|---|---|---|---|
| T3 | 43584548 | 6.5 | 98.18 | 0.02 | 98.15 | 94.63 | 50.85 |
| T4 | 44004608 | 6.6 | 98.15 | 0.02 | 98.23 | 94.85 | 51.33 |
| T8 | 44110978 | 6.6 | 98.64 | 0.02 | 98.06 | 94.35 | 49.39 |
| T11 | 41209658 | 6.2 | 98.17 | 0.03 | 97.95 | 94.10 | 49.57 |
| T17 | 45164800 | 6.8 | 98.10 | 0.03 | 97.93 | 94.05 | 48.96 |
| T20 | 41158010 | 6.2 | 98.23 | 0.02 | 98.10 | 94.47 | 49.60 |
| T24 | 42413318 | 6.4 | 98.19 | 0.02 | 98.06 | 94.34 | 49.73 |
| T32 | 46182444 | 6.9 | 97.02 | 0.02 | 98.12 | 94.56 | 49.92 |

A. Library Preparation and Sequencing
1 Sample Quality Control
2 Library Construction, Quality Control and Sequencing
B. Results and Instructions
1 Data Quality Control
1.1 Distribution of Sequencing Quality
1.2 Distribution of Sequencing Error Rate
1.3 Distribution of A/T/G/C Base
1.4 Results of Raw Data Filtering
2 Summary of Sequencing Data Information
C. Appendix
1 Introduction of Sequencing Data Format
2 Explanation of Sequencing Data Related
3 References

| | | | | | | | |
|---|---|---|---|---|---|---|---|
| T35 | 41252534 | 6.2 | 97.82 | 0.03 | 97.88 | 93.93 | 50.35 |
| T36 | 49439348 | 7.4 | 97.14 | 0.02 | 98.10 | 94.42 | 47.22 |
| T38 | 44719236 | 6.7 | 98.00 | 0.02 | 98.11 | 94.49 | 48.72 |
| T39 | 51810798 | 7.8 | 98.17 | 0.02 | 98.01 | 94.27 | 49.85 |
| T40 | 41603472 | 6.2 | 97.87 | 0.02 | 98.20 | 94.76 | 51.17 |
| T42 | 43906992 | 6.6 | 98.01 | 0.02 | 98.25 | 94.82 | 49.74 |
| T43 | 39726814 | 6.0 | 98.14 | 0.02 | 98.04 | 94.35 | 51.12 |
| T44 | 43762324 | 6.6 | 98.24 | 0.02 | 98.11 | 94.33 | 49.14 |
| T45 | 43007870 | 6.5 | 97.58 | 0.02 | 98.07 | 94.30 | 48.46 |
| T48 | 56537514 | 8.5 | 98.13 | 0.02 | 98.09 | 94.37 | 49.29 |
| D49 | 46689628 | 7.0 | 98.40 | 0.02 | 98.23 | 94.80 | 51.50 |
| D50 | 51573528 | 7.7 | 96.99 | 0.02 | 98.16 | 94.62 | 47.83 |
| D54 | 45618906 | 6.8 | 97.89 | 0.02 | 98.28 | 94.95 | 50.49 |
| D55 | 50059592 | 7.5 | 98.24 | 0.02 | 98.06 | 94.43 | 51.66 |
| D56 | 46132804 | 6.9 | 97.76 | 0.02 | 98.21 | 94.68 | 50.21 |
| D58 | 46295328 | 6.9 | 97.74 | 0.02 | 98.01 | 94.27 | 50.24 |
| D59 | 46813260 | 7.0 | 98.07 | 0.03 | 97.95 | 94.09 | 51.44 |
| D62 | 42865624 | 6.4 | 98.17 | 0.02 | 98.13 | 94.62 | 51.66 |
| D64 | 44570244 | 6.7 | 98.08 | 0.03 | 97.87 | 93.83 | 50.01 |
| D65 | 52676698 | 7.9 | 98.26 | 0.02 | 98.13 | 94.50 | 50.91 |
| D66 | 44000934 | 6.6 | 97.98 | 0.02 | 98.07 | 94.27 | 50.29 |
| D67 | 40489444 | 6.1 | 98.13 | 0.02 | 98.10 | 94.48 | 51.55 |
| D68 | 49037312 | 7.4 | 97.16 | 0.02 | 98.23 | 94.86 | 50.17 |
| D70 | 39249794 | 5.9 | 97.27 | 0.02 | 98.23 | 94.85 | 50.74 |
| D72 | 45763236 | 6.9 | 97.69 | 0.02 | 98.12 | 94.57 | 50.47 |
| D73 | 46906792 | 7.0 | 97.99 | 0.02 | 98.20 | 94.73 | 51.01 |
| D74 | 48820782 | 7.3 | 98.39 | 0.02 | 98.07 | 94.23 | 50.08 |
| D77 | 48091806 | 7.2 | 98.34 | 0.02 | 98.00 | 94.22 | 51.68 |
| D78 | 45947632 | 6.9 | 98.01 | 0.02 | 98.14 | 94.60 | 50.43 |
| D80 | 42179218 | 6.3 | 97.85 | 0.02 | 98.23 | 94.77 | 50.83 |
| T37 | 39638018 | 5.9 | 96.59 | 0.02 | 98.25 | 95.04 | 52.29 |
| T1 | 40249880 | 6.0 | 97.29 | 0.03 | 97.15 | 92.60 | 51.88 |
| T2 | 49274884 | 7.4 | 97.51 | 0.03 | 97.65 | 93.65 | 50.77 |
| T5 | 40182512 | 6.0 | 97.15 | 0.03 | 97.00 | 92.32 | 50.44 |
| T6 | 49018944 | 7.4 | 97.12 | 0.03 | 97.62 | 93.54 | 50.79 |
| T7 | 54589032 | 8.2 | 97.41 | 0.03 | 97.12 | 92.60 | 51.86 |
| T10 | 61492096 | 9.2 | 97.70 | 0.03 | 96.93 | 92.12 | 51.45 |
| T12 | 41451202 | 6.2 | 95.73 | 0.03 | 97.16 | 92.68 | 50.91 |
| T13 | 46585738 | 7.0 | 97.23 | 0.03 | 97.26 | 92.80 | 52.04 |
| T14 | 45088894 | 6.8 | 96.85 | 0.03 | 96.92 | 92.14 | 51.26 |
| T15 | 40983132 | 6.1 | 97.38 | 0.03 | 96.88 | 92.06 | 50.74 |
| T16 | 44598826 | 6.7 | 97.04 | 0.03 | 97.00 | 92.24 | 52.12 |
| T18 | 39687584 | 6.0 | 97.43 | 0.03 | 96.92 | 92.12 | 50.89 |
| T19 | 52161296 | 7.8 | 97.67 | 0.03 | 97.07 | 92.43 | 51.87 |
| T21 | 51750950 | 7.8 | 97.11 | 0.03 | 96.95 | 92.23 | 50.95 |
| T22 | 52053472 | 7.8 | 97.48 | 0.03 | 97.14 | 92.54 | 51.75 |
| T23 | 45927170 | 6.9 | 97.51 | 0.03 | 96.96 | 92.22 | 50.42 |
| T25 | 43504528 | 6.5 | 97.34 | 0.03 | 97.07 | 92.42 | 52.17 |
| T26 | 42102112 | 6.3 | 96.95 | 0.03 | 96.98 | 92.22 | 50.86 |
| T27 | 42807780 | 6.4 | 97.18 | 0.03 | 96.77 | 91.76 | 50.44 |
| T28 | 46527946 | 7.0 | 96.86 | 0.03 | 97.36 | 93.00 | 52.21 |
| T29 | 44216802 | 6.6 | 96.94 | 0.03 | 97.21 | 92.73 | 51.23 |
| T30 | 46821972 | 7.0 | 96.99 | 0.03 | 97.34 | 92.99 | 51.09 |

A. Library Preparation and Sequencing
1 Sample Quality Control
2 Library Construction, Quality Control and Sequencing
B. Results and Instructions
1 Data Quality Control
1.1 Distribution of Sequencing Quality
1.2 Distribution of Sequencing Error Rate
1.3 Distribution of A/T/G/C Base
1.4 Results of Raw Data Filtering
2 Summary of Sequencing Data Information
C. Appendix
1 Introduction of Sequencing Data Format
2 Explanation of Sequencing Data Related
3 References

| | | | | | | | |
|---|---|---|---|---|---|---|---|
| T34 | 52662376 | 7.9 | 96.68 | 0.03 | 97.30 | 92.93 | 51.65 |
| T41 | 44782916 | 6.7 | 96.63 | 0.03 | 96.90 | 92.04 | 50.20 |
| T46 | 39840150 | 6.0 | 96.95 | 0.03 | 96.81 | 91.88 | 51.46 |
| T47 | 46362534 | 7.0 | 96.83 | 0.03 | 97.35 | 93.00 | 50.23 |
| D51 | 46462332 | 7.0 | 96.97 | 0.03 | 97.28 | 92.97 | 51.34 |
| D52 | 44550058 | 6.7 | 96.87 | 0.03 | 96.46 | 91.11 | 50.71 |
| D53 | 50494132 | 7.6 | 96.98 | 0.03 | 97.42 | 93.18 | 51.73 |
| D57 | 66073860 | 9.9 | 97.18 | 0.03 | 96.93 | 92.16 | 51.26 |
| D60 | 47828914 | 7.2 | 97.08 | 0.03 | 97.25 | 92.84 | 49.86 |
| D61 | 44290794 | 6.6 | 96.82 | 0.03 | 97.17 | 92.63 | 51.69 |
| D63 | 46759066 | 7.0 | 96.66 | 0.03 | 97.27 | 92.78 | 50.97 |
| D69 | 49000504 | 7.4 | 96.81 | 0.03 | 96.94 | 92.20 | 51.78 |
| D71 | 50767464 | 7.6 | 95.78 | 0.03 | 96.91 | 92.10 | 50.78 |
| D75 | 40357932 | 6.1 | 95.67 | 0.03 | 97.09 | 92.50 | 51.17 |
| D76 | 49885418 | 7.5 | 95.40 | 0.03 | 97.47 | 93.25 | 50.73 |
| D79 | 43612294 | 6.5 | 95.89 | 0.03 | 97.19 | 92.72 | 51.39 |
| T9 | 55083296 | 8.3 | 97.19 | 0.03 | 97.10 | 92.56 | 51.41 |

Sample: sample name
Raw reads: total amount of reads of raw data, each four lines taken as one unit. For paired-end sequencing, it equals the amount of read1 and read2, otherwise it equals the amount of read1 for single-end sequencing.
Raw data: (Raw reads) * (sequence length), calculating in G. For paired-end sequencing like PE150, sequencing length equals 150, otherwise it equals 50 for sequencing like SE50.
Effective: (Clean reads/Raw reads)*100%
Error: base error rate
Q20, Q30: (Base count of Phred value > 20 or 30) / (Total base count)
GC: (G & C base count) / (Total base count)



### C. Appendix

#### 1 Introduction of Sequencing Data Format

The original raw data from Illumina platform are transformed to Sequenced Reads, known as Raw Data or RAW Reads, by base calling. Raw data are recorded in a FASTQ file, which contains sequencing reads and corresponding sequencing quality. Every read in FASTQ format is stored in four lines, as indicated below (Cock P.J.A. et al. 2010):

```
@HWI-ST1276:71:C1162ACXX:1:1101:1208:2458 1:N:0:CGATGT
NAAGAACACGTTCGGTCACCTCAGCACACTTGTGAATGTCATGGGATCCAT
+
#55???BBBBB?BA@DEEFFCFFHHFFCFFHHHHHHHHFAE0ECFFD/AEHH
```

Line 1 begins with a '@' character and is followed by the Illumina Sequence Identifiers and an optional description. The details of Illumina sequence identifier are listed in the table below.

| Identifier | Meaning |
|---|---|
| HWI-ST1276 | Instrument – unique identifier of the sequencer |
| 71 | Run number – Run number on instrument |
| C1162ACXX | Flow Cell ID – ID of flow cell |
| 1 | Lane Number – positive integer |
| 1101 | Tile Number – positive integer |
| 1208 | X – x coordinate of the spot. Integer which can be negative |
| 2458 | Y – y coordinate of the spot. Integer which can be negative |
| 1 | Read number. 1 can be single read or Read 2 of paired-end. |
| N | Y if the read is filtered (did not pass), N otherwise. |
| 0 | Control number - 0 when none of the control bits are on, otherwise it is an even number |
| CGATGT | Illumina index sequences |

A. Library Preparation and Sequencing
1 Sample Quality Control
2 Library Construction, Quality Control and Sequencing
B. Results and Instructions
1 Data Quality Control
1.1 Distribution of Sequencing Quality
1.2 Distribution of Sequencing Error Rate
1.3 Distribution of A/T/G/C Base
1.4 Results of Raw Data Filtering
2 Summary of Sequencing Data Information
C. Appendix
1 Introduction of Sequencing Data Format
2 Explanation of Sequencing Data Related
3 References

Line 3 begins with a '+' character and is optionally followed by the same sequence identifiers and descriptions as in Line 1.

Line 4 encodes the quality values for the bases in Line 2 and contains the same number of characters as the bases in the read (Cock, 2009.).

#### 2 Explanation of Sequencing Data Related

(1) The data delivered is a compressed file in format of '.fq.gz'. Before data delivery, we will calculate the md5 value of each compressed file and please check it when you get the data. There are two ways to check the md5 value. In Linux environment, you can use 'md5sum -c <*md5.txt>' command under the data directory. In Windows environment, you can use a calibration tool e.g. hashmyfiles. If the md5 value of compressed file doesn't match with the one we provide in md5 file in data directory, the file may have been damaged during the transmitting procedure.

(2) For paired-end (PE) sequencing, every sample should have 2 data flies (read1 file and read2 file). These 2 files have the same line number, you could use 'wc -l' command to check the line number in Linux environment. The line number divide by 4 is the number of reads.

(3) The data size is the space occupied by the data in the hard disk. It's related to the format of the disk and compression ratio. And it has no influence on the quantity of sequenced bases. Thus, the size of read1 file may be unequal to the size of read2 file.

(4) When customer’s samples need large amount of data e.g. whole genome sequencing data, we would use separate-lane sequencing strategy to make sure the quality of data, so it's possible that one sample has several parts sequencing data. For example, if sample 1 has two read1 files, sample1_L1_1.fq.gz and sample1_L2_1.fq.gz, that means this sample was sequenced on different lanes.

(5) If we agree to deliver the clean data before the project starts, we will filter the data strictly according to the standard to obtain high quality clean data which can be used for further research and paper writing. We will discard the paired reads in the following situation: when either one read contains adapter contamination; when either one read contains more than 10 percent uncertain nucleotides; when either one read contains more than 50 percent low quality nucleotides (base quality less than 5). The data analysis results based on the clean data that is filtered by this standard can be approved by high level magazines (Yan L.Y. et al . 2013). If you want to get more information, please refer to the official website of Novogene (www.novogene.com).

(6) The Index is normally in the middle of the adapter during the process of experiment and sequencing except the special library. We can get the Read1 sequence and Read2 sequence according to the Index read. They are all the sequence of samples so that it's no necessary to dispose the beginning and end of reads in the downstream analysis (e.g. mapping).

(7) 30 days after the data delivery, we will delete outdated data. So please keep your data properly. If you have any questions or concerns, please contact us as soon as possible.
